# ProtXAI: Explainable AI Reveals Structural Determinants of Protein Dynamics

**DOI:** 10.64898/2026.05.26.727866

**Authors:** Faraneh Haddadi, Joan Planas-Iglesias, Jan Mican, Jana Horackova, Sérgio M. Marques, Matej Demovic, Pavel Kohout, Jiri Damborsky, David Bednar, Stanislav Mazurenko

**Author notes:** author for correspondence: Stanislav Mazurenko.

## Abstract

Molecular dynamics simulations provide atomistic views of protein motions, but conventional analyses often struggle with extracting subtle mechanistic insights from complex trajectories. Here, we present an integrated framework, ProtXAI, combining molecular dynamics and explainable artificial intelligence (XAI), to identify residue-level determinants of conformational change across diverse protein systems. By leveraging inter-residue distance dynamics, deep learning, and sequential relevance propagation, the approach captures both local fluctuations and long-range communication pathways within protein structures. We applied this framework to three mechanistically distinct systems: apolipoprotein E4 (ApoE4), staphylokinase (SAK) variants, and an ancestral luciferase. Across these applications, our XAI-based approach recovered experimentally supported dynamic hotspots: ligand-responsive hinges in ApoE4, mutation-dependent flexibility shifts in SAK, and evolutionary redistribution of motions in the luciferase. ProtXAI also revealed additional long-range couplings not accessible to classical analysis. Together, these findings demonstrate that combining molecular dynamics with XAI provides a general and scalable strategy for dissecting protein dynamics and uncovering structural determinants of function, stability, and evolutionary changes without prior bias. This approach thus advances the current methodological repertoire for analysing proteins and their intrinsic properties.

**Highlights:** Molecular dynamics simulations are increasingly accessible, yet scalable tools for comparative analysis remain limited.

We demonstrate that machine learning coupled with explainable AI can automatically extract structural determinants of protein dynamics from trajectories.

ProtXAI identifies key dynamic regions across diverse scenarios, including comparison of protein variants, understanding ligand modulation, and single-trajectory analysis.

ProtXAI enables scalable, unbiased interpretation of long trajectories, providing an alternative to manual, time-intensive analysis.

## 1. Introduction

Molecular dynamics (MD) simulations provide atomistic, time-resolved insights into protein structure and dynamics. Despite their successes in explaining conformational landscapes, ligand-binding mechanisms, folding pathways, and ion transport processes, such simulations often still require visual inspection and expert curation, for example, to identify appropriate collective variables (CVs)^1,2^. Traditional computational methods for analysing MD trajectories, such as RMSD, RMSF, and principal component analysis (PCA), often struggle to capture subtle, high-dimensional conformational changes and distributed collective motions^3,4^. These approaches are frequently subjective and are designed and optimized separately for each protein system, relying on manually selected structural observables rather than a standardized, automated framework^5–7^. As the scale and complexity of MD datasets continue to increase, there is a growing need for automated, scalable, and unbiased analysis strategies^8^.

Machine learning (ML) has become a powerful complement to MD simulations, enabling the identification of nonlinear structural patterns, classification of functional states, and efficient analysis of large trajectory ensembles ^9,10^. Beyond trajectory analysis, ML is now widely used to construct neural network–based force fields and potential energy surfaces with near *ab initio* accuracy, including Behler–Parrinello neural network potentials and ANI models^11–13^. More recently, generative and diffusion-based approaches such as AlphaFold3 and BioEmu, together with ML-assisted enhanced sampling methods, have enabled direct exploration of protein conformational ensembles beyond conventional MD timescales^14–16^. Despite these advances, limited interpretability of many ML architectures remains a major obstacle to mechanistic insight, particularly for high-dimensional structural data^17,18^. Accordingly, explainable artificial intelligence (XAI) approaches have emerged to map model predictions back to physically meaningful molecular features such as residue interactions and collective motions^17^. The utility of ML for extracting structural determinants directly from MD trajectories has been demonstrated in studies identifying binding determinants in SARS-CoV spike protein complexes and detecting allosteric communication pathways from simulation data^19–21^.

While impactful, current approaches for using ML to identify critical protein elements have several limitations. They often focus on a specific selected structural element, e.g., predefined interfaces or specific amino acids of the protein^19^. They also usually treat MD frames as a set of independent observations, thus relying on the identification of the patterns in ensembles of static structures rather than their dynamical motions^20^. In addition, many existing approaches rely on relatively coarse attribution strategies, such as feature importance derived from linear models^22^ or gradient-based saliency maps, which often provide residue-level or global importance scores without resolving pairwise interactions or propagating relevance through deep network layers in a mechanistically consistent manner^17,18^.

In this work, we build on these efforts and propose a more expressive explainability framework based on layer-wise relevance propagation (LRP), enabling fine-grained, residue-level attribution across deep models, which we termed ProtXAI (Figure 1). Rather than restricting the analysis to known binding sites, ProtXAI systematically evaluates all residues to capture potential allosteric communication and long-range couplings, which are increasingly recognized as central to protein function^23^. We evaluated ProtXAI across three representative protein engineering scenarios: (i) ligand-induced stabilization of apolipoprotein E4 (ApoE4), simulated with and without drug candidate 3-sulfopropanoic acid (3-SPA), reflecting small-molecule modulation of conformational equilibria; (ii) discrimination of staphylokinase (SAK) variants within a conserved fold, modelling sequence-driven redistribution of flexibility; and (iii) intrinsic dynamic coordination along an evolutionary trajectory of a Renilla luciferase, capturing how functional optimization reshapes collective motions. Together, these scenarios span key protein engineering challenges, including protein stabilization, mutational redesign, and functional evolution. Comparison of the resulting residues identified by ProtXAI with what we learned in our previous work revealed that ProtXAI looked beyond the simple flexibility analysis and managed to capture allosterically important residues determining the differences in protein dynamics.

**Figure 1.**
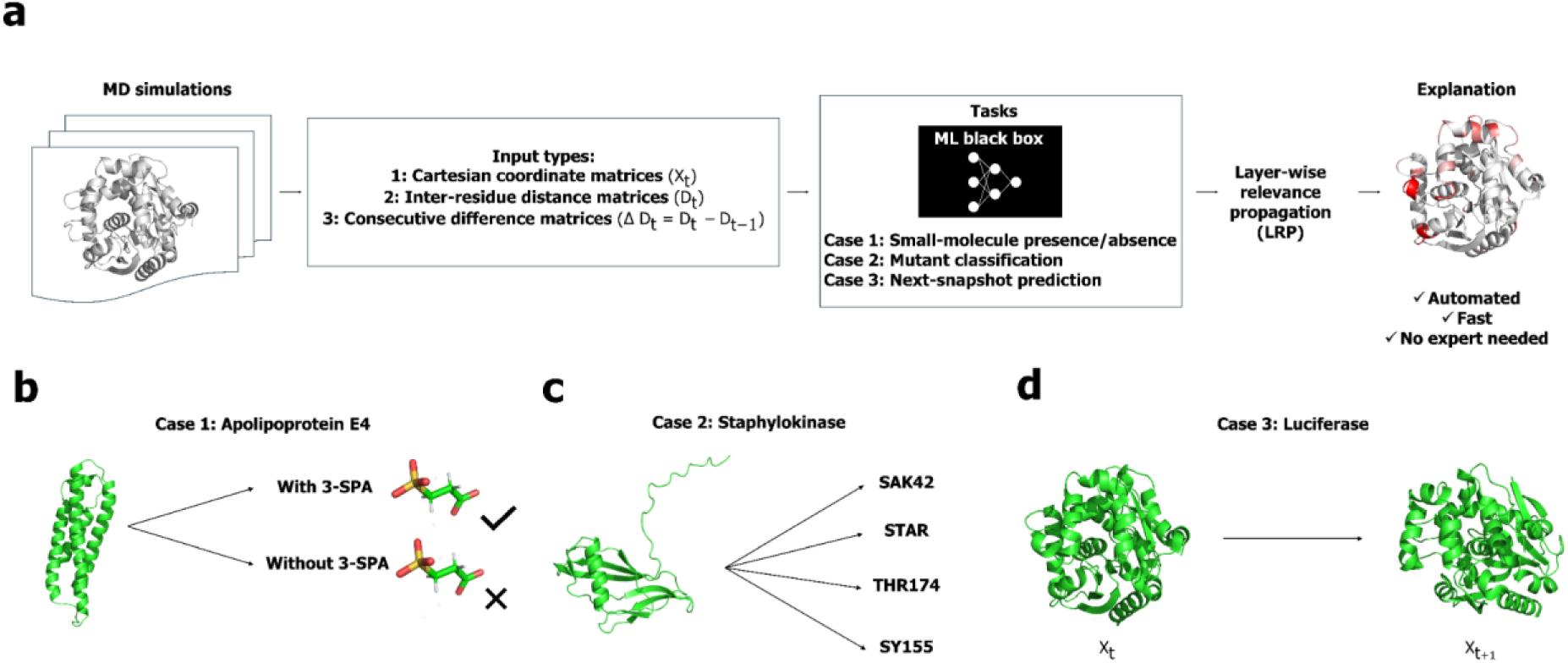
Overview of the ProtXAI pipeline integrating MD simulations with ML and XAI. (a) General workflow. MD trajectories are first converted into structural representations, including Cartesian coordinate matrices (Xₜ), inter-residue distance matrices (Dₜ), and frame-to-frame distance-difference matrices (ΔDₜ = Dₜ − Dₜ₋₁). These representations are used as inputs to supervised ML models tailored to each biological question. Model outputs are subsequently analysed using layer-wise relevance propagation (LRP) to obtain residue- and residue-pair–level relevance maps, enabling mechanistic interpretation of protein dynamics without manual feature selection. (b) ApoE4 drug candidate case: identification of residue interactions and dynamic regions that differ between the apo state and the small-molecule–bound state (3-sulfopropanoic acid, 3-SPA), revealing ligand-induced redistribution of motion. (c) SAK variants case: characterization of mutation-dependent dynamic differences among engineered variants (SAK42D, STAR, THR174, and SY155), identifying structural hotspots underlying variant-specific flexibility and stability. (d) Luciferase evolutionary case: detection of residues coordinating intrinsic conformational transitions along an evolutionary trajectory, highlighting how redistribution of collective motions accompanies functional optimization.

## 2. Materials and Methods

### 2.1 Overview of the ProtXAI Pipeline

ProtXAI is a general workflow that transforms MD trajectories into residue-level mechanistic interpretations through supervised ML and XAI. The framework consists of four main stages:

#### Step 1. Data preprocessing and representation

MD trajectories are subsampled and coarse-grained to Cα atoms to capture backbone conformational dynamics. Pairwise residue–residue distances are first computed for each frame, and distance-difference matrices (ΔDₜ = Dₜ − Dₜ₋₁) are subsequently derived to capture short-timescale dynamical changes in residue–residue interactions. For coordinate-based modelling tasks, Cartesian coordinates of Cα atoms extracted from aligned trajectories are used as inputs. The resulting datasets are partitioned into training and test sets using system-specific splitting strategies designed to minimize information leakage (see below).

#### Step 2. Supervised model training

A convolutional neural network (CNN) is trained to address a task-specific objective, such as state discrimination or next-frame prediction. Distance-based representations are modelled using two-dimensional CNN layers to capture spatial correlations between residue pairs, whereas coordinate-based representations are modelled using one-dimensional CNN architectures. The networks consist of convolutional layers followed by fully connected layers for classification tasks or encoder–decoder architectures for next-frame prediction. Model performance is evaluated on an independent test set.

#### Step 3. Explainability via LRP

Following training, LRP is applied to redistribute the model’s output backward through the network. This assigns a relevance value to each input feature for every individual MD frame, quantifying the contribution of specific residues or residue pairs to the model’s decision.

#### Step 4. Relevance aggregation and residue-level interpretation

Frame-level relevance values are aggregated across time and, when applicable, across spatial dimensions (e.g., averaging over x/y/z coordinates or symmetrizing residue-pair matrices) to obtain stable per-residue or residue-pair relevance profiles. These aggregated relevance maps identify dynamic hotspots and long-range couplings underlying the modelled behaviour.

### 2.2. MD Simulations of Model Systems

#### ApoE4 with or without 3-SPA^24^

Full-length ApoE4 was generated by homology modelling using ApoE3 (PDB ID: 2L7B^25^) as a template. Systems were protonated at pH 7.4 using PROPKA^26^, solvated in mTIP3P water^27^, ion-neutralized to 0.1 M salt concentration, and parameterized using the CHARMM36m^28^ force field. After minimization and equilibration steps, production simulations were performed for ApoE4 in the apo state (the absence of ligand) and in the presence of 100 molecules of 3-SPA distributed around the protein. Adaptive sampling^29^ was conducted using the root-mean-square deviation (RMSD)^30^ as sampling metric and one-dimensional time-lagged independent component analysis (tICA)^31^ over 20 epochs, yielding an aggregate simulation time of 20 μs. Snapshots were subsampled every 5 ns and restricted to residues 24–161, resulting in 8,036 MD frames. To minimize information leakage arising from adaptive sampling, data were split using an epoch-based 80/20 partitioning scheme. The first 10% of frames within each test epoch were excluded to reduce temporal correlation between training and test sets, yielding 6,560 training and 1,476 test samples. System-specific representations and model architectures followed the general ProtXAI workflow described above.

#### SAK variants^32^

SAK variants (SAK42D, STAR, SY155, and THR174) were modelled using AlphaFold3 and protonated using H++^33^. Crystallographic waters from PDB ID: 1BUI^34^ were incorporated where applicable. Systems were prepared using the AMBER ff19SB^35^ force field and OPC water model^36^, neutralized, minimized, equilibrated, and subjected to production MD simulations. Each variant was simulated for 200 ns per replica (three replicas per variant; 600 ns total per variant). Snapshots were extracted every 0.1 ns, yielding 23,956 frames in total across all variants. Classification was formulated as a four-class problem distinguishing the variants. An 80/20 random split was applied, resulting in 19,165 training and 4,791 test samples. Data representations and model configurations followed the general pipeline described in Section 2.1.

#### Luciferase subfamily^37^

MD simulations were performed for RLuc8 (PDB ID: 2PSF, chain B)^38^, Anc^HLD-Rluc^ (PDB ID: 6G75)^39^, and AncFT (PDB ID: 6S97)^37^. Systems were protonated using H++^33^, solvated in TIP3P water^40^ with ≥10 Å padding, and neutralized to 0.1 M ionic strength. Because trajectory data for AncFT were not available from the original study, this system was re-simulated under consistent preparation and adaptive sampling protocols. Trajectories were recorded every 0.1 ns, aligned to a reference structure to remove translational and rotational drift, and subsampled every 5 ns. Cartesian coordinates of Cα atoms were extracted for downstream modelling. The task was formulated as a self-supervised next-frame prediction problem, in which each MD snapshot served as input and the subsequent snapshot as the target. Data were randomly split into 80% training and 20% test subsets.

### 2.3. Data Extraction and Representation

MD trajectories were aligned to a reference structure to remove global translational and rotational motions, ensuring that subsequent analysis reflects only internal conformational changes. To reduce temporal redundancy and computational cost, frames were subsampled at regular intervals prior to feature extraction.

Protein structures were represented using Cα atoms to provide a compact description of backbone dynamics. Pairwise residue–residue distances were first computed for each frame using the PyEMMA software package, resulting in distance matrices (Dₜ) that describe the instantaneous structural state of the protein. To capture dynamical information, distance-difference matrices were subsequently constructed as the change in pairwise Euclidean distances between consecutive frames (ΔDₜ = Dₜ − Dₜ₋₁). This representation encodes short-timescale dynamical variations in residue–residue interactions, enabling the model to focus on temporal changes rather than static structural features. The use of distance-difference features was motivated by preliminary experiments showing that static distance-based representations could be trivially classified by simple models and were of limited mechanistic value, whereas capturing dynamical changes requires more expressive architectures and better reflects the underlying physical processes.

For supervised learning tasks (ApoE4 and SAK systems), input samples consisted of distance-difference matrices labelled according to the corresponding condition, such as ligand presence or protein variant identity. For the luciferase system, a self-supervised formulation was employed, in which each MD frame was used to predict the subsequent frame using the same representation, enabling the model to learn intrinsic conformational dynamics without explicit labels. Datasets were partitioned into training, validation, and test sets using system-specific strategies designed to minimize bias (see MD Simulations of Model Systems). To prevent temporal leakage, consecutive frames were not assigned to different subsets.

### 2.4. Supervised Machine Learning

Following the data preparation described above, different ML strategies were used depending on the task. For ApoE4 and SAK, which relied on distance-difference matrices, 2D CNNs were used to capture spatial correlations. For the Luciferase systems, which used coordinate-level data, a 1D CNN-based autoencoder was used to predict the next MD snapshot. A detailed summary of model architectures, data representations, and training configurations for all systems is provided in Supplementary Tables S1 and S2.

#### ApoE4: Binary Classifier to Distinguish ApoE4 With and Without Drug Candidate 3-SPA

A supervised 2-D CNN classifier was trained to distinguish apo (unbound) versus 3-SPA–bound ApoE4 using the distance-difference representation. The network architecture consisted of two convolutional layers with 4 and 8 filters, followed by two fully connected layers with 16 and 32 neurons. ReLU activations were applied throughout, except for the final sigmoid output layer. To reduce overfitting observed during preliminary experiments, we used a dropout rate of 0.4 together with L2 regularization (λ = 0.1). As a baseline comparison, a logistic regression model with ElasticNet regularization was also trained on the same distance-difference inputs.

#### SAK: Multi-Class Classification of Four Protein Systems

A four-class CNN classifier was trained to distinguish SAK42D, STAR, SY155, and THR174. The architecture included two 2D convolutional layers (2 and 4 filters), a flattening layer, and a dense layer with 8 neurons. The output layer used a Softmax activation across four classes. ReLU activations, L2 regularization (λ = 0.1), and dropout (0.05) were applied. A multinomial logistic regression baseline with ElasticNet regularization (C = 0.01, l1_ratio = 0.5) using the SAGA solver served as a comparison.

#### Luciferase: Autoencoder for Next-Frame Prediction

The luciferase systems were modelled using a 1D CNN autoencoder that predicts the next MD snapshot from the current one. The encoder consisted of two layers with 16 and 8 neurons and a 3-neuron bottleneck. The decoder mirrored this architecture. All layers used ReLU activations. Training used mean squared error loss between predicted and actual coordinates. The same architecture was used for all luciferase proteins.

### 2.5. XAI Analysis

XAI techniques were applied to all trained models using LRP via the iNNvestigate library^41^. LRP redistributes the prediction backwards through the network to assign relevance scores to each input feature. (1) For ApoE4 and SAK, LRP produced symmetric 2-D relevance maps with the same dimensions as the distance matrices, enabling residue-pair–level interpretation. (2) For luciferase, relevance values had the same shape as the coordinate input; relevance was averaged across x/y/z for each residue and then aggregated across snapshots to obtain a single stable relevance score per residue. We used **neuron_selection_mode=’all’** to propagate relevance across all outputs simultaneously. Across all models, the **lrp.sequential_preset_a** rule was applied for robustness and stability during propagation.

### 2.5 ANM calculations

Structures of RLuc (PDB ID: 2psf, chain B^38^), Anc^HLD-RLuc^ (PDB ID: 6g75, chain A^39^), and AncFT (PDB ID: 6s97, chain A^37^) were used as input to ProDy^42^ implementation of ANM to obtain prediction of B-factors and the analysis of predicted cross-correlations along the different residues in the proteins. All parameters were used as set by default, except the number of normal modes that was set to the length of the protein sequence minus one, as per the software manual recommendations. Predicted B-factors are herein reported as obtained from the software, while the reported cross-correlations correspond to the average of the absolute cross-correlation value of a residue *i* to any other residue *j*.

### 2.6 Shortest Path Map calculations

The Shortest Path Map (SPM) was computed from three MD replicas following the protocol described previously^43^. For each replica, trajectories were aligned to a representative structure identified as the frame with the lowest RMSD to the replica-averaged structure. The distance and correlation matrices were calculated using whole MD trajectories. The SPM calculations were performed using the dedicated web server available at http://spmosuna.com4242. We used the default calculation settings (Significance: 0.3; Distance: 6 Å).

## 3. Results and Discussion

### 3.1. Method development

We developed ProtXAI as a general framework for extracting mechanistic insights from MD trajectories using supervised ML and XAI. The pipeline integrates structural representations derived from MD simulations with neural network models and *post hoc* interpretability analysis (Figure 1a). MD trajectories are converted into numerical representations of protein motion, including residue–residue distance matrices, distance-difference matrices capturing short-timescale dynamical changes, and Cartesian coordinates of Cα atoms. These representations serve as inputs to CNNs trained on task-specific objectives, after which XAI methods are used to map model decisions back to structurally meaningful residues and residue interactions. The overall goal of this workflow is to automatically identify dynamic regions and residue interactions that differentiate functional states or coordinate conformational transitions.

To evaluate the robustness of the ML component, we tested multiple model configurations and training strategies across the three systems. These included variations in network architecture, hyperparameters (learning rate, regularization strength, and dropout), and data representations capturing either structural states or short-timescale dynamical changes (Tables S1 and S2). Model performance was assessed using held-out test sets and cross-validation procedures appropriate for each system, with classification accuracy and reconstruction error used as evaluation metrics depending on the task. Across all cases, the selected architectures achieved stable performance and generalization, indicating that the models successfully captured informative dynamical patterns from the MD data.

To interpret the trained models and relate their predictions to physical protein features, we applied XAI analysis based on LRP. LRP redistributes the model output backwards through the network, assigning relevance scores to input features while preserving conservation of the prediction score across layers. Compared with gradient-based saliency approaches, LRP provides more stable and physically interpretable attribution for deep neural networks applied to structured inputs such as residue-distance matrices. In preliminary benchmarking experiments performed within our group, LRP consistently produced clearer and more robust residue-level relevance maps than alternative attribution methods, motivating its use in the ProtXAI pipeline. In addition to its interpretability, the approach is computationally efficient after model training, allowing rapid generation of residue-level relevance maps across large MD ensembles and making it suitable for scalable comparative analysis of protein dynamics. For each of our case studies, the typical time to extract a single set of XAI relevance values from an MD simulation was around 10 minutes on a regular laptop (CPU), suggesting the possibility of systematically processing large libraries of protein variants.

### 3.2. Method validation using model proteins

We next applied this framework to three representative protein engineering scenarios. In the first case, we analysed apolipoprotein E4 (ApoE4) in the presence and absence of a drug candidate, the small molecule 3-sulfopropanoic acid (3-SPA), to identify residue interactions and dynamic regions associated with ligand-induced stabilization. In the second case, we compared four engineered staphylokinase (SAK) variants to characterize mutation-dependent redistribution of conformational flexibility within a conserved fold. In the third case, we investigated the luciferase evolutionary series using a self-supervised learning formulation in which the model predicts subsequent MD frames, enabling identification of residues that coordinate intrinsic conformational transitions. Together, these case studies illustrate how ProtXAI can reveal dynamic hotspots and long-range couplings across diverse protein systems.

#### Small molecule effects on the dynamics of ApoE4

ApoE4 is a central lipid transport protein and a major genetic risk factor for Alzheimer’s disease. Small molecules, such as 3-SPA, have been shown to stabilize ApoE4 by modulating its conformational dynamics, yet the structural interactions underlying this stabilization are not fully resolved^24,44^. While hydrogen–deuterium exchange mass spectrometry (HDX-MS) and MD simulations demonstrate ligand-induced changes at a global level, they do not directly reveal which residue–residue interactions are the most informative for distinguishing the ligand-bound state from the apo form.

To address this, we combined adaptive-sampling MD simulations with an explainable machine-learning framework trained on changes in residue-pair distances within the ApoE4 N-terminal domain. A 2D CNN was trained to classify whether a given pair of MD snapshots contains ApoE4 in the presence of 3-SPA or not. The relevance analysis was used to identify the residue pairs (Figure 2a) and regions (Figure 2c) most responsible for the model’s decisions. Before applying the CNN, we evaluated whether a simpler baseline model could solve the task. The baseline logistic regression model performed close to a random predictor (test accuracy 0.45, F1 = 0.60; Table S3), while the 2D CNN maintained strong predictive performance (test accuracy 0.95, F1 = 0.95). This benchmarking confirmed that the dynamic representation captures non-linear patterns in residue motion that cannot be recovered by a simple linear model, providing a more appropriate basis for subsequent XAI analysis.

**Figure 2.**
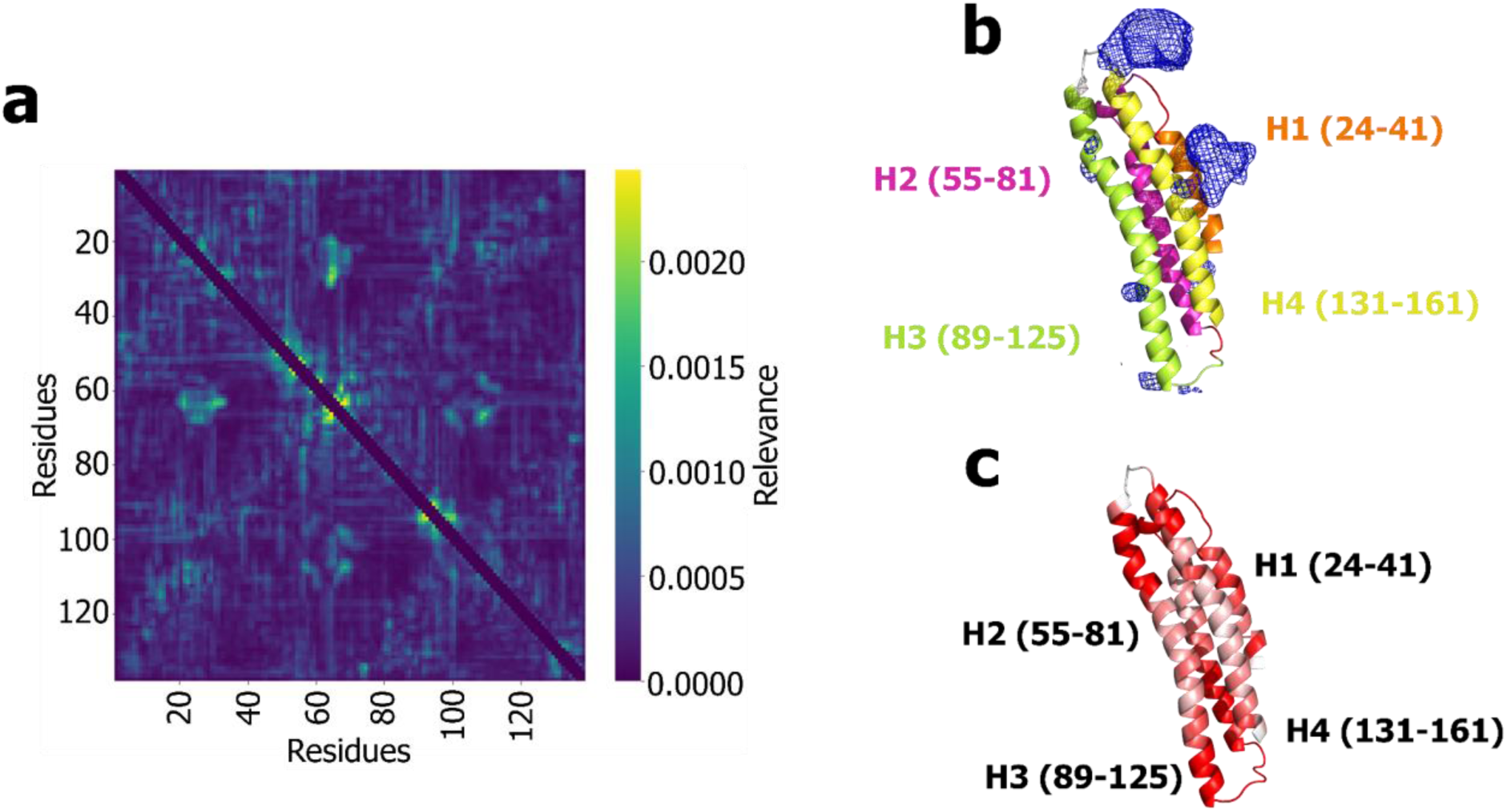
XAI analysis of the ApoE4 system in the presence and absence of 3-SPA. (a) Heatmap showing relevance scores assigned to residue–residue distance features by LRP. Higher values indicate residue pairs that contribute most strongly to distinguishing the ligand-bound and apo states. Relevance values were averaged across all analysed frames. (b) Spatial distribution of ligand-associated relevance mapped onto the ApoE4 structure obtained from MD simulations^43^. Blue isosurfaces represent regions with high ligand density as visualized in VMD (Visual Molecular Dynamics). The structure is shown as a cartoon representation with major α-helices labelled H1–H4 (H1: orange, H2: magentas, H3: green, and H4: yellow). The numbers indicate the residue ranges corresponding to each helix: H1 (24–41), H2 (55–81), H3 (89–125), and H4 (131–161). (c) Average relevance score per residue projected onto the ApoE4 structure, highlighting dynamic regions most affected by the presence of 3-SPA. Together, these analyses reveal that ligand-associated dynamical changes localize primarily to the H2–H3 hinge and adjacent helices, indicating that stabilization by 3-SPA is mediated through modulation of motion in these helical regions rather than through a single localized interaction site.

The XAI analysis identified a small set of residue pairs with consistently high relevance, most prominently interactions involving residues 87, 90, and 91, as well as interactions with residues 66, 81, and 117 (Figure 2b). When mapped onto the structure, these signals localize to the hinge between helices H2 and H3. Interestingly, additional contributions to high relevance values were observed at both ends of helix H3 and at the beginning of helix H4, which were not usually in contact with the ligand. This organization indicates that the model captured both direct ligand-sensitive regions and longer-range effects on ApoE4 dynamics.

We recently investigated this system using hydrogen–deuterium HDX-MS and MD simulations combined with classical analysis approaches^44^. Without requiring prior knowledge or manual inspection of MD trajectories, ProtXAI identified the same key regions previously reported. In particular, HDX-MS and MD experiments showed that the H2–H3 hinge (residues 68–97) and the H3–H4 junction (residues 113–135) undergo major dynamical changes upon 3-SPA binding. These regions were recovered with high relevance scores in our XAI analysis (Figure 2c).

Notably, previous work demonstrated that 3-SPA interacts with charged residues at the ApoE4 self-association interface and induces a shift toward an ApoE3-like conformational ensemble, accompanied by stabilization and partial straightening of helix H3^44^. These effects propagate through long-range structural coupling originating from the C112R mutation, which disrupts key inter-residue interactions and reorganizes the global fold. The strong agreement between these experimentally and computationally characterized regions and the XAI-derived relevance hotspots indicates that the model captures mechanistically important interaction networks underlying ligand-induced stabilization rather than merely local flexibility.

At the same time, XAI identified additional regions, such as residues 26–27 and 34–54, that did not show strong ligand-dependent stabilization in HDX-MS. These signals likely reflect long-range allosteric couplings captured by the distance-difference representation rather than local solvent exposure or unfolding. This interpretation is consistent with previous findings that structural perturbations in ApoE4 propagate over long distances through a domino-like mechanism, linking distal regions of the protein through coordinated changes in residue interaction^44^.

To further characterize these ligand-induced effects, we compared residue-level relevance scores with B-factors obtained from simulations with and without 3-SPA (Figure S1). While linear correlations were moderate in the ligand-bound state and weaker in its absence, cosine similarity remained high in both cases (0.81 and 0.76, respectively), indicating that relevance captured similar residue-level patterns of flexibility. This suggests that the model identified dynamic regions consistent with experimentally and computationally observed flexibility, with stronger agreement under ligand-bound conditions.

#### Mutational effects on the dynamics of SAK

SAK is a bacterial plasminogen activator considered an alternative thrombolytic agent, but its clinical application is limited by its immunogenicity^45,46^. This has motivated several studies engineering multiple SAK variants to preserve fibrinolytic function while reducing immune recognition through a small number of targeted mutations^32^. Although the resulting variants (SAK42D, STAR, SY155, and THR174) share the same overall fold, experimental studies revealed clear differences in stability, aggregation propensity, and flexibility. These observations imply that the introduced sequence changes primarily perturb protein dynamics rather than static structure. Thus, pinpointing the exact regions across the four variants that are responsible for these perturbations can shed light on the mechanisms behind the observed differences in experimental measurements.

Available experimental and simulation techniques for SAK42D, STAR, SY155, and THR174, including HDX-MS, NMR ensemble variability and relaxation, crystal B-factors, ML-based flexibility predictors, classical MD, and adaptive-sampling MD28, consistently indicated that variant-dependent behaviour is localized within a small set of structural regions. These include the β2–H1 hinge (residues 49–56), the H1–β3 loop, the β3–β4 loop (82–86), and the C-terminal β6 strand (Figure S2), which repeatedly emerge as the principal dynamic hotspots distinguishing the variants.

To assess whether these experimentally observed differences can be recovered and interpreted directly from MD data, we applied the ProtXAI pipeline trained on residue-pair distance changes. This approach tests whether XAI can identify the same dynamic regions highlighted experimentally and provide a mechanistic link between conformational motion and measured biophysical behaviour. As for the ApoE4 system, we first evaluated whether a simple baseline model could capture the variant-specific dynamical signatures. The logistic regression baseline performed no better than a random predictor (test accuracy 0.17, F1 = 0.17; Table S4), whereas the 2-D CNN achieved near-perfect performance (test accuracy 0.99, F1 = 0.99). This comparison demonstrates that the distance-difference representation captures dynamic information that cannot be extracted by linear models and provides a suitable foundation for interpreting variant-specific dynamical features using XAI.

The XAI analysis reproduced all major experimentally supported hotspots across the four variants. Particularly, the H1–β3 region (residues 58–77), which is a hallmark of the non-immunogenic variants SY155 and THR174, was strongly captured both by the residue-level relevance profiles (Figure 3d) and by the highest-scoring residue-pair interactions (Figure 3a, Table S4).

**Figure 3.**
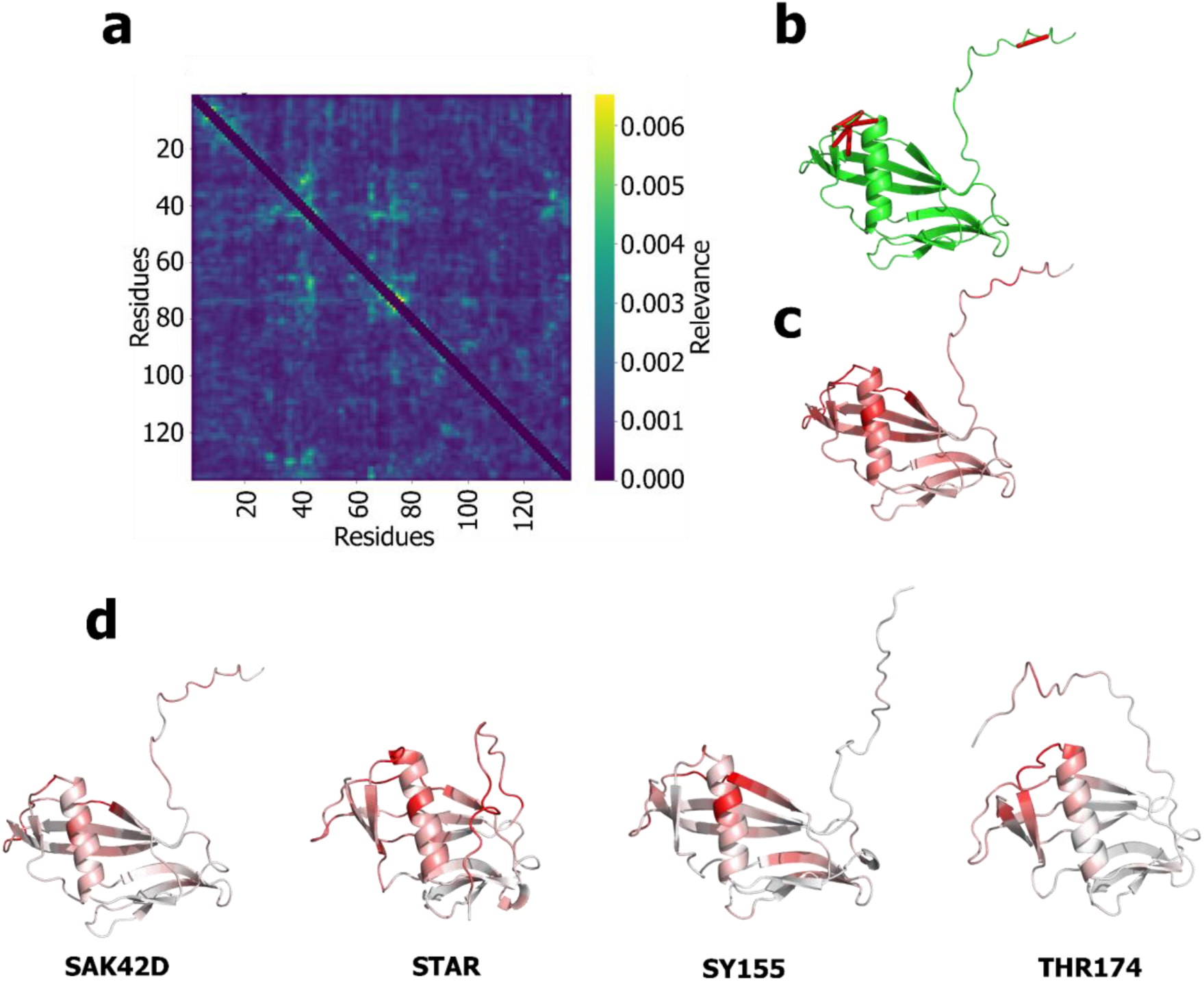
Relevance analysis of the SAK variants. (a) Heatmap showing residue-pair relevance values derived from layer-wise relevance propagation and averaged across all four SAK variants. Higher values indicate residue interactions that contribute most strongly to distinguishing the variants. (b) The five residue pairs with the highest relevance scores are mapped onto the SAK42D structure and connected by red lines (see Table S5 for details). (c) Residue-level relevance projected onto the SAK42D structure. A single relevance value per residue was obtained by averaging each row of the residue-pair relevance matrix shown in panel (a). Red indicates higher relevance values. (d) Average residue-level relevance scores projected onto the structures of the four SAK variants (SAK42D, STAR, SY155, and THR174), highlighting variant-specific patterns of dynamical importance. Together, these analyses identify a small set of interconnected dynamic hotspots, most prominently the β2–H1 hinge region (Figure S2) and adjacent loop elements, that differentiate the variants and explain experimentally observed differences in flexibility and stability within the conserved fold.

The XAI analysis reproduced all major experimentally supported hotspots across all four variants, including SAK42D^31^ (Figure 3d). The surrounding β2–H1 hinge consistently emerged as a universal dynamic hub. Additional agreement was observed at the β3–β4 loop, the C-terminal end of helix H1, and the β6 strand, with variant-specific relevance patterns that closely mirror trends observed in HDX-MS, NMR, and MD. In SAK42D, relevance was primarily concentrated in the N-terminal region and across the β1–β2 segment (residues ∼35–44), extending toward the β2–H1 hinge, with additional contributions from the central portion of helix H1 and the H1–β3 loop, reflecting localized baseline dynamics at the β-sheet–helix interface. In STAR, relevance extended across a broader set of regions, including the N-terminus (residues ∼3–21), the β1–β2 segment (residues ∼33–45), the central region of helix H1, and the H1–β3 loop extending into the β3 strand (residues ∼71–83), as well as the C-terminal β6 strand, indicating a more distributed coupling between the N-terminus, helix H1, and downstream β-structures. In SY155, relevance remained cantered around the β1–β2 region (residues ∼35–45) and the central portion of helix H1, with additional contributions from the H1–β3 loop (residues ∼72–74) and distal β-strands including β4 and β6, consistent with experimentally observed flexibility increases in these regions. In THR174, relevance was strongly concentrated in the H1–β3 region (residues ∼69–78), with additional contributions from the N-terminus, the β1–β2 loop, and the β6 strand, in line with experimental observations identifying this variant as the most dynamically perturbed, exhibiting elevated flexibility in helix H1 and the adjacent β3 region.

Beyond these overlaps, XAI revealed long-range couplings that were weak or absent in experimental readouts. These include pronounced N-terminal involvement in STAR and THR174, as well as internal interaction networks in SY155, particularly involving residue pairs linking the H1–β3 region (∼65–75) with β1–β2 (∼35–45) and adjacent secondary structure elements. These patterns suggested the presence of coordinated, long-range communication pathways captured by distance-based representations rather than by local unfolding or exchange processes alone. Motivated by these findings, we applied the Shortest Path Method (SPM) to further investigate allosteric pathways (Figure S4). The analysis indicated that the allosteric path traversed regions identified by XAI, located in close proximity to the H1–β3 loop, particularly in STAR, supporting their role in mediating long-range coupling. However, SPM alone did not differentiate between variants, as the inferred pathways remain largely conserved across them. In contrast, XAI resolved distinct, variant-specific coupling patterns by enforcing discrimination among all four variants, highlighting differences in how these pathways are utilized rather than their overall topology.

Taken together, this integrated analysis showed that differences among SAK variants arise from the redistribution of motion across a small number of interconnected structural elements. XAI not only confirmed experimentally established dynamic hotspots but also uncovered hidden communication pathways, particularly clusters of relevant interactions between residues ∼65–75 that distinguish non-immunogenic variants from the WT. Importantly, the functional relevance of the H1–β3 loop is not solely determined by its structural proximity to the protein core, but also by its positioning along communication pathways that traverse this region. The high-relevance interactions identified by XAI lie in close spatial proximity to SPM-derived pathways, suggesting that they reflect underlying signal propagation routes. While SPM revealed largely conserved pathways and therefore lacks discriminative power, XAI captured subtle but functionally meaningful differences in dynamical organization, particularly distinguishing non-immunogenic variants from STAR, thereby providing a more detailed view of how minimal sequence changes reshape the dynamical landscape.

### Determinants of the luciferase dynamics

Renilla-type luciferases and their reconstructed ancestors provide a well-established model for studying how protein dynamics shape enzymatic function and evolution^28^. The ancestral scaffold Anc^HLD-Rluc^ is thermostable and bifunctional but weakly luminescent, whereas engineered descendants AncFT and the modern luciferase RLuc8 exhibit markedly enhanced catalytic efficiency and altered bioluminescence behaviour. Extensive experimental work showed that these functional gains arise not from changes in the catalytic core, but from redistribution of conformational dynamics within the cap domain, particularly involving Loop 9 and the α4–α5 region^37^. This evolutionary series, therefore, offers a suitable benchmark for evaluating whether explainable machine-learning methods applied to MD simulations can recover known dynamic determinants and track changes in motion along an evolutionary trajectory.

In Anc^HLD-Rluc^, we analysed raw Cartesian coordinates from MD simulations using a two-dimensional CNN-based autoencoder combined with explainability analysis to identify residues most influential for predicting conformational changes. Loop 9 emerged as the dominant relevance hotspot (Figure 4), consistent with its experimentally and computationally established flexibility^37^. Additional relevance peaks were detected in structurally constrained regions with low B-factors, indicating residues that may contribute to coordinating global motions rather than exhibiting local flexibility alone.

**Figure 4.**
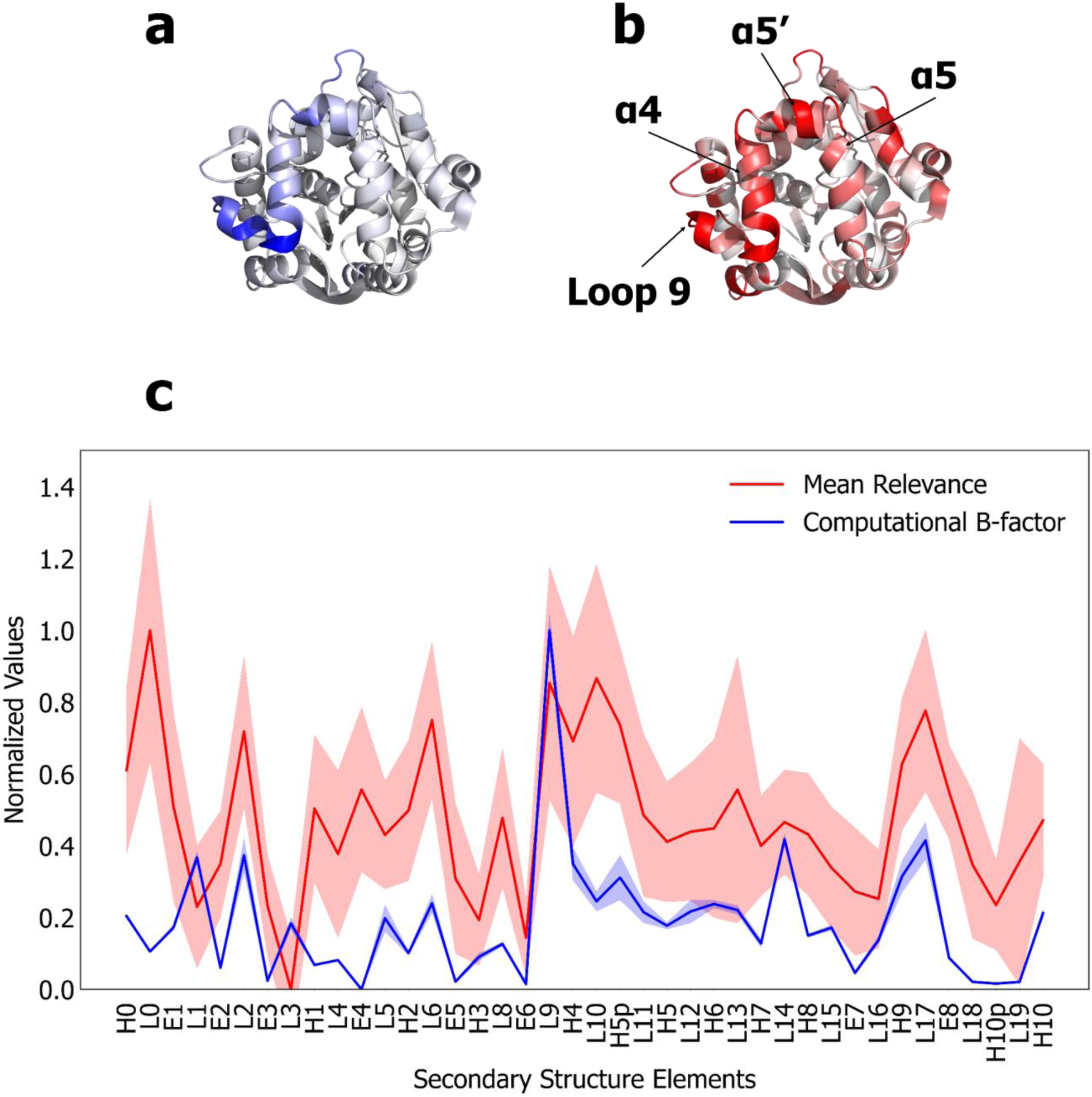
Overview of structural flexibility and XAI analysis of Anc^HLD-Rluc^. (a) B-factor values mapped onto the protein structure. (b) Results of the XAI pipeline projected onto the structure. (c) Comparative plot of computational B-factor values (blue) calculated from the full MD trajectories and mean relevance scores (red) across secondary structure elements. Both profiles were normalized to the [0,1] range after aggregation to enable direct comparison. Shaded regions represent the standard error of the mean (SEM) calculated across independent relevance runs and independent adaptive sampling trajectories used for B-factor estimation.

To quantify how the XAI signal related to experimentally observed dynamics, we compared residue-level relevance scores with computational B-factors and HDX-MS measurements aggregated across secondary structure elements. As summarized in Figure S5, the normalized datasets show moderate-to-strong agreement between the three descriptors. Pearson correlations reach r = 0.50 between relevance and B-factors and r = 0.66 between relevance and HDX-MS, while cosine similarities indicate strong pattern overlap (cos θ = 0.71 and 0.94, respectively). Spearman correlations (ρ = 0.51 and ρ = 0.62) further confirm that regions with elevated experimental flexibility tend to receive high relevance scores. These results show that the XAI-derived importance maps recover key functional regions identified experimentally, particularly the L9–α4 fragment, including Loop 9 (residues 140–157). The strong LRP signal observed at residues 147–152 and 155–158 aligns with Loop 9, while residues 160–168 map to the α4 helix, confirming that the model identified this region as a major dynamical feature.

We next compared the relevance signal with predictions from an anisotropic network model (ANM) to evaluate whether the learned patterns correspond to collective motions in the protein fold. At the residue level, relevance correlated moderately with B-factors (ρ ≈ 0.42) and with ANM amplitudes (ρ ≈ 0.52), indicating that residues identified by the neural network often correspond to dynamically active regions participating in collective motions rather than merely reflecting local flexibility (Figure S6). Restricting the analysis to the structured region of the protein (residues 38–155) strengthened these relationships, yielding a higher correlation between relevance and B-factors (ρ = 0.54) and ANM (ρ = 0.78), respectively (Figure S7). Aggregating values at the secondary-structure level produced similar trends (Figure S8), supporting the interpretation that XAI highlighted residues that coordinate collective dynamics within the fold.

To further probe how these residues participate in long-range communication pathways, we analysed the protein using a Shortest Path Map method^43^, identifying residue chains that connect dynamically relevant regions across the structure. The contributions of relevance, ANM amplitudes, and B-factors were quantified for each path (Table S6), revealing that several chains associated with Loop 9 and adjacent cap-domain elements account for a substantial fraction of the total relevance signal. The residues forming these communication paths (Table S7) overlap strongly with experimentally identified dynamical regions, particularly around Loop 9 and the α4–α5 segment. Correlation analysis using the shortest-path representation (Figure S9) showed a weak positive association between relevance and B-factors (Pearson r = 0.14; Spearman ρ = 0.19), accompanied by high cosine similarity, indicating strong agreement in their pathway-level patterns despite limited linear correlation.

In contrast, relevance exhibits a moderate positive correlation with ANM amplitudes (Pearson r = 0.46; Spearman ρ = 0.51), indicating that residues identified as important by the model tend to participate in collective motions captured by low-frequency normal modes. These results further support the conclusion that the XAI model captures residues associated with the propagation of conformational changes. Notably, several SPM-identified residues, particularly within SPM path_10 (140–150, 153, 158–159), coincide with the L9–α4 region and strongly overlap with the LRP-derived importance peaks. Additional agreement was observed in regions around residues 81–90, 165, 204, and 223, providing consistent evidence that these regions are recurrently identified as important in the protein dynamics in the protein.

To place these findings in an evolutionary context, we applied the same XAI workflow to AncFT and RLuc8, whose dynamic behaviour has been extensively characterized. Previous experimental and computational studies demonstrated a progressive shift in the dominant dynamics from Loop 9 toward the α4–α5 region along the evolutionary path leading to improved catalytic performance^37^. Our XAI analysis reproduced this redistribution: while Anc^HLD-Rluc^ retained strong relevance in Loop 9, AncFT and RLuc8 showed increased relevance along the α4–α5 region, consistent with stabilization of helix α4 and reorganization of cap-domain motion (Figure S10).

Finally, we compared the relevance distribution with an independent structural metric describing the spatial extent of residue motion (Sphere_Scale), which reflects how widely individual residues explore their local conformational space. While direct linear and monotonic correlations between Sphere_Scale and relevance values were negligible (Spearman ρ = 0.06, Pearson r = 0.001), the cosine similarity between the two profiles was high (cos θ = 0.92), indicating that both descriptors highlight similar residue-level spatial patterns despite substantial differences in their relative magnitudes and scaling (Figure S11). In comparison, Sphere_Scale and B-factors showed weak and slightly negative correlations (Spearman ρ = −0.18, Pearson r = −0.11) together with only moderate cosine similarity (cos θ = 0.55), suggesting limited overall agreement between these descriptors. Relevance values and B-factors, however, exhibited moderate positive correlations (Spearman ρ = 0.41, Pearson r = 0.49) and relatively high cosine similarity (cos θ = 0.68), indicating that the XAI-derived relevance maps capture experimentally supported flexibility trends while also reflecting additional structural features related to collective conformational dynamics rather than merely identifying the most flexible residues.

## 4. Conclusions

In this study, we investigated whether combining supervised or self-supervised learning with XAI can help identify critical elements governing protein dynamics. Although recent studies have demonstrated the interpretability of latent representations in protein language models and autoencoder-based frameworks through sparse or adversarial architectures, these approaches primarily focus on feature extraction from learned embeddings or latent spaces rather than directly identifying residue-level determinants of molecular dynamics behaviour. In contrast, our developed pipeline ProtXAI operates directly on MD-derived structural representations and uses XAI to localize physically meaningful residues and couplings associated with conformational dynamics, ligand binding, and mutational effects. Furthermore, unlike traditional CV-based analyses that require predefined reaction coordinates or dimensionality-reduction assumptions, ProtXAI does not rely on manually selected CV. Instead, the framework learns discriminative dynamic patterns directly from simulation data while retaining residue-level interpretability.

We evaluated ProtXAI in three representative scenarios: ligand-induced stabilization of ApoE4 in the presence or absence of drug candidate 3-SPA^32,44^, mutation-dependent redistribution of conformational flexibility among engineered SAK variants, and intrinsic protein dynamics analysed through a self-supervised next-frame prediction task in selected luciferases^37^. Across all three systems, integrating MD simulations with XAI enabled the identification of structural determinants underlying ligand effects, mutational changes, and evolutionary shifts in protein dynamics.

In ApoE4, the analysis revealed stabilization of hinge regions and redistribution of motion upon 3-SPA binding. In SAK variants, differences in experimental stability and flexibility were traced to a small set of interconnected dynamic hotspots within the conserved fold. In luciferases, intrinsic dynamics were governed by residues coordinating global motions rather than simply exhibiting high local flexibility. ProtXAI recovered experimentally validated hotspots, including the SPA-responsive H2–H3 and H3–H4 hinges in ApoE4, the β2–H1 hinge and β6 strand in SAK, and Loop 9 in luciferases, while also identifying additional long-range couplings not directly accessible to experimental techniques.

While showing significant promise, our pipeline represents a first step in extracting the residue-level relevance from MD trajectories using XAI, and several limitations warrant future research. Our three case studies represent typical protein engineering scenarios for leveraging MD data, yet our analysis is far from being systematic as the selection of proteins was primarily driven by the available expertise in the group and knowledge of a particular protein system. Larger MD datasets annotated on the residue level would enable more thorough benchmarking and eventually accelerate the development of such pipelines but are currently missing. Although a definitive ground truth for residue-level importance in protein dynamics is difficult to define, future validation through mutagenesis, perturbation simulations, and experimental measurements such as HDX-MS^47^ or NMR^3^ will further provide annotations. Another interesting direction to explore is the granularity of the relevance attribution. We briefly explored this direction by aggregating the relevance on the secondary structure level. More thorough investigation of different aggregation methods or even ML-derived coarse-graining^48^ can be very attractive, for example, in loop grafting^49^.

Despite these limitations, the consistency of the identified relevance patterns across three mechanistically distinct systems demonstrates that ProtXAI captures biologically meaningful dynamical information rather than simple artifacts. Importantly, the framework not only recovers experimentally established dynamic hotspots but also reveals coordinated long-range couplings and communication pathways that are difficult to identify using conventional analysis approaches alone. Since ProtXAI operates directly on MD-derived structural representations without requiring predefined collective variables or manually selected descriptors, it provides a scalable and largely unbiased strategy for interpreting increasingly large molecular simulation datasets. More broadly, this study highlights how interpretable ML can bridge the gap between high-dimensional simulation data and mechanistic understanding, opening new opportunities for studying allostery, conformational regulation, ligand modulation, and protein engineering at residue-level resolution.

## Acknowledgements

This work was supported by the European Union’s Horizon 2020 research and innovation programme under grant agreements No. 857560 (CETOCOEN) and Horizon Europe Framework programme No. 101136607 (CLARA). Computational resources were provided by the e-INFRA CZ, ELIXIR-CZ, and RECETOX RI projects (90254, LM2023055, and LM2023069), supported by the Ministry of Education, Youth and Sports of the Czech Republic. This publication reflects only the author’s view, and the European Commission is not responsible for any use that may be made of the information it contains. Brno Ph.D. Talent Scholarship holders Pavel Kohout and Jan Mičan acknowledge funding from the Brno City Municipality.

## Code availability

The code and necessary scripts to reproduce the results will be made publicly available on GitHub upon publication.

## Author Contributions

F.H. designed and performed the machine learning and explainable AI analyses, processed molecular dynamics data, and wrote the manuscript. J.P.-I., J.M., J.H., S.M.M., M.D., and P.K. contributed to molecular dynamics simulations, data preparation, and interpretation of results.

J.D. and D.B. supervised the research and contributed to conceptualization and manuscript revision. S.M. conceived the study, supervised the project, and contributed to methodology development and writing. All authors reviewed and approved the final manuscript.

---

**Figure S1.**
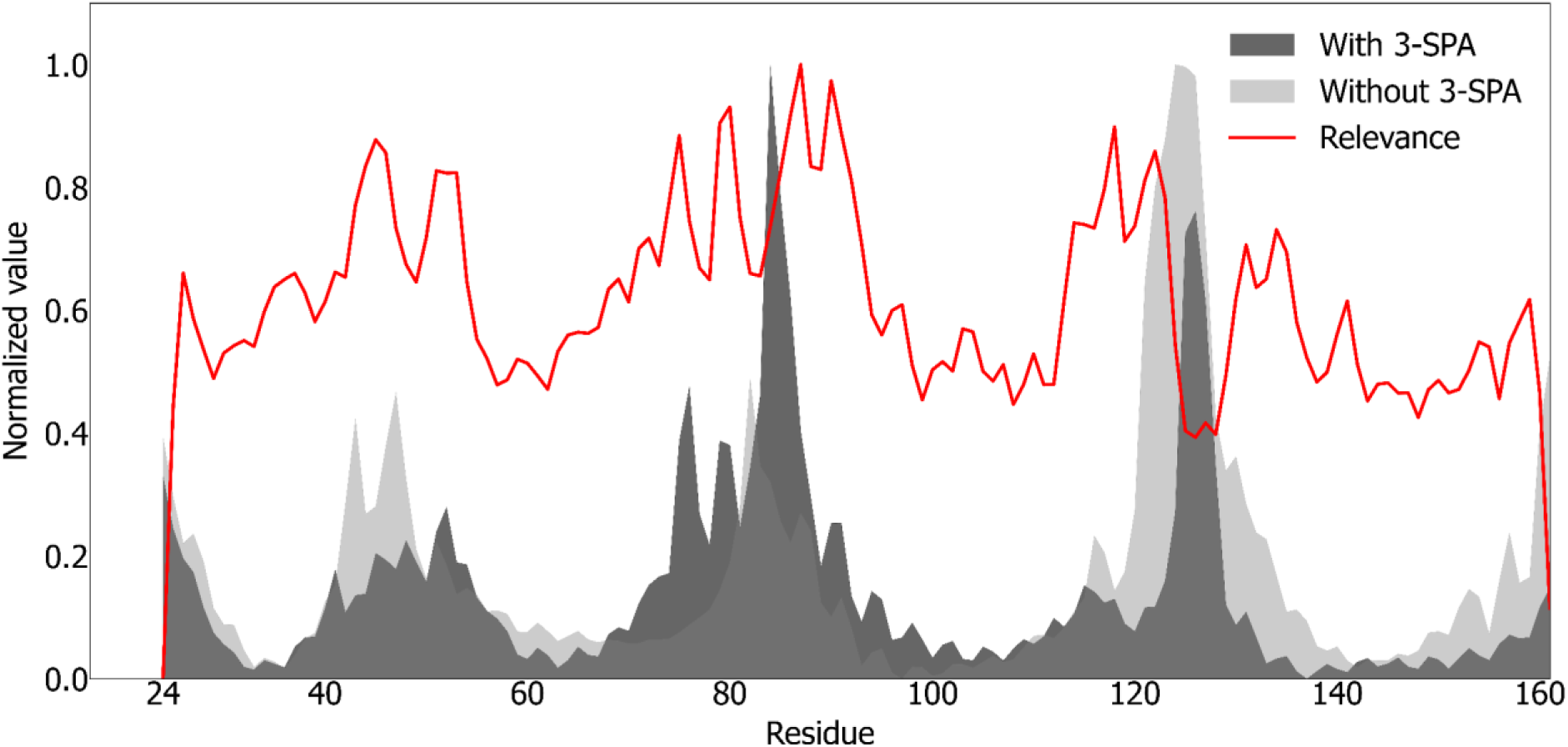
Correlation analysis between LRP-derived relevance scores and B-factors for ApoE4 in the presence and absence of 3-SPA. Relevance shows moderate agreement with B-factors in the presence of 3-SPA (Pearson 0.27, Spearman 0.48) and weaker linear correlation in its absence (Pearson 0.05, Spearman 0.30), while cosine similarity remains high in both cases (0.81 and 0.76, respectively). These results indicate that relevance captures similar residue-level patterns of flexibility, particularly in the ligand-bound state, despite differences in absolute scaling.

**Figure S2.**
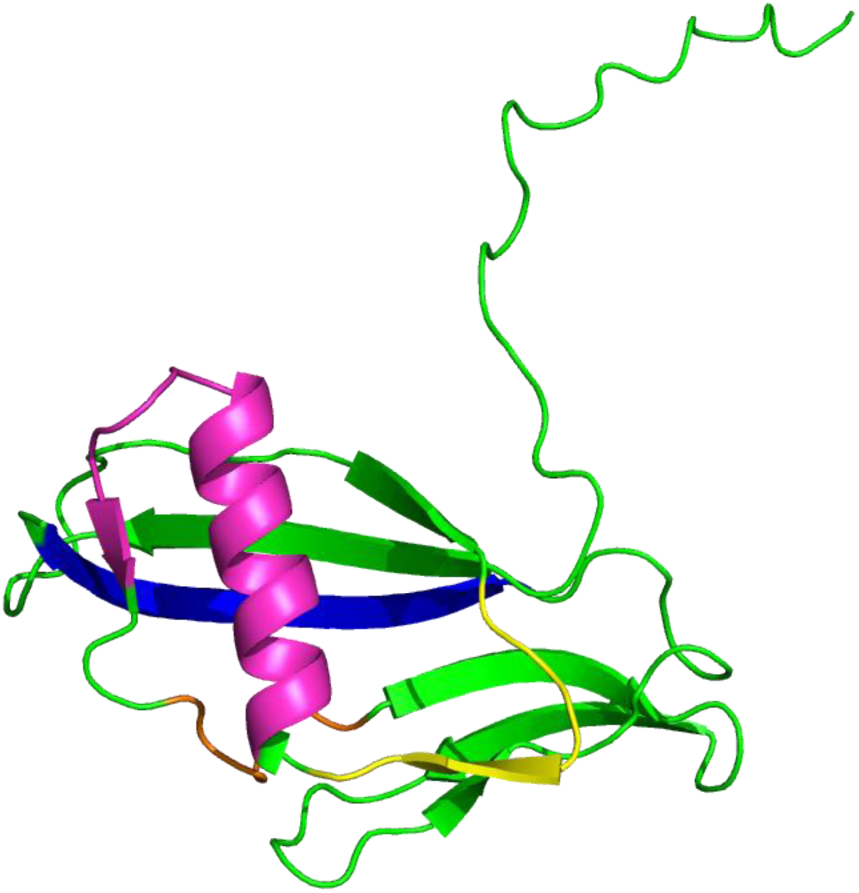
Structural mapping of key dynamic regions. Cartoon representation of the protein highlighting regions identified as principal dynamic hotspots. The β2–H1 hinge (residues 49–56) is shown in yellow, the H1–β3 region (residues 58–77) in magenta, the β3–β4 loop (residues 82–86) in orange, and the C-terminal β6 strand in blue. The remaining structure is depicted in green. These regions consistently emerge as major contributors to the dynamic differences between variants.

**Figure S3.**
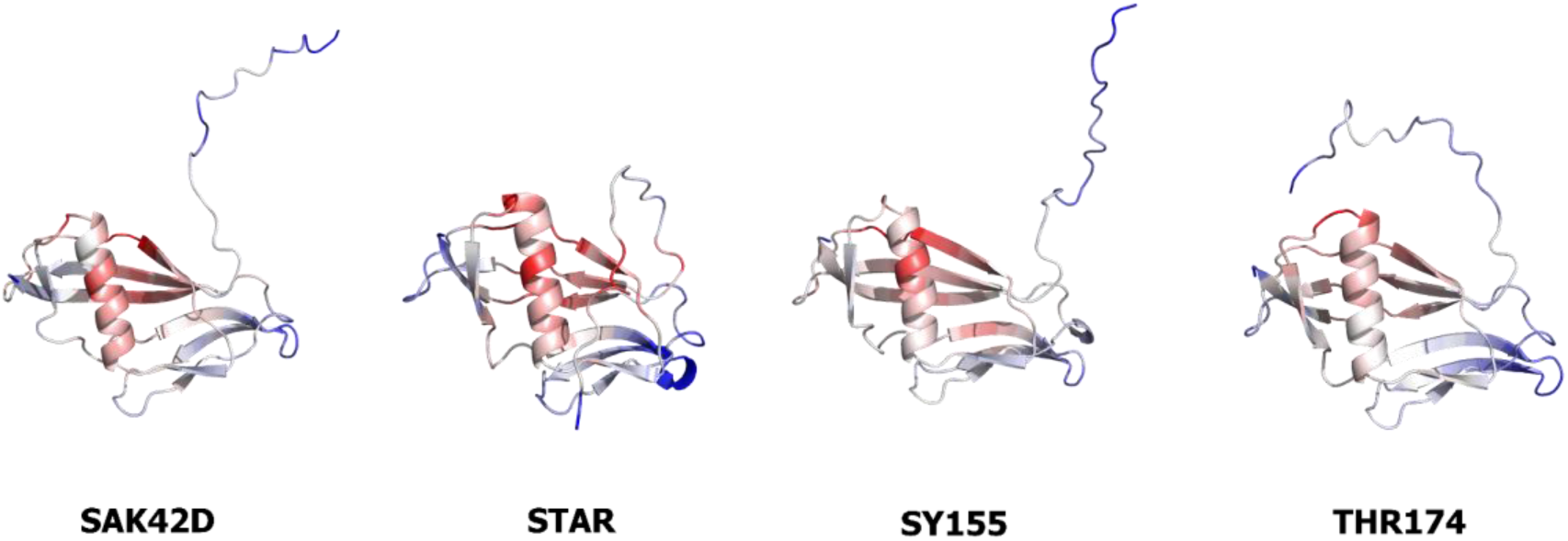
Comparison of model-derived relevance and structural flexibility across SAK variants. Structures are coloured according to the difference between relevance values and B-factors: red highlights residues with higher relevance (key features identified by the model), whereas blue indicates regions with higher B-factors, corresponding to increased atomic mobility.

**Figure S4.**
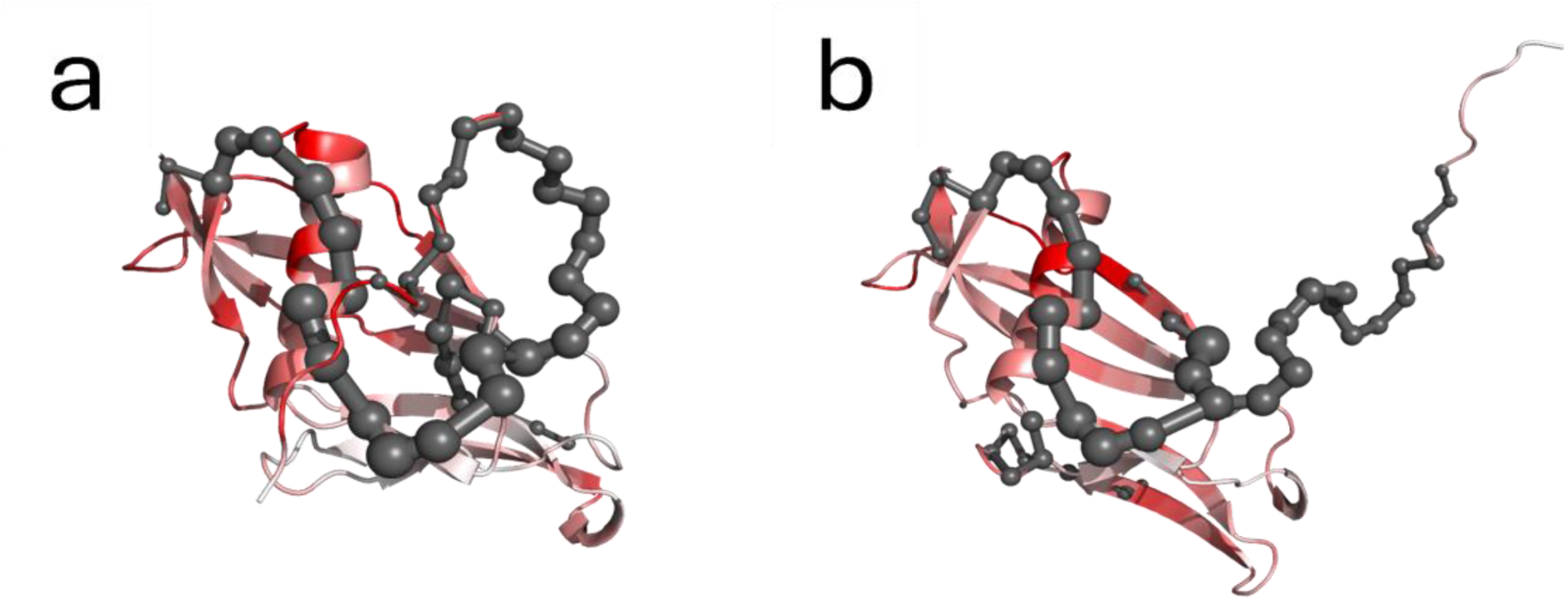
Shortest Path Map annotation for A: STAR and B: SY155 protein variants, in addition to the relevance scores in red. The SPM highlights the allosteric pathways traversing through relevance-annotated residues. Although SPM covers similar residues for both variants, the XAI picks different relevance-related regions.

**Figure S5.**
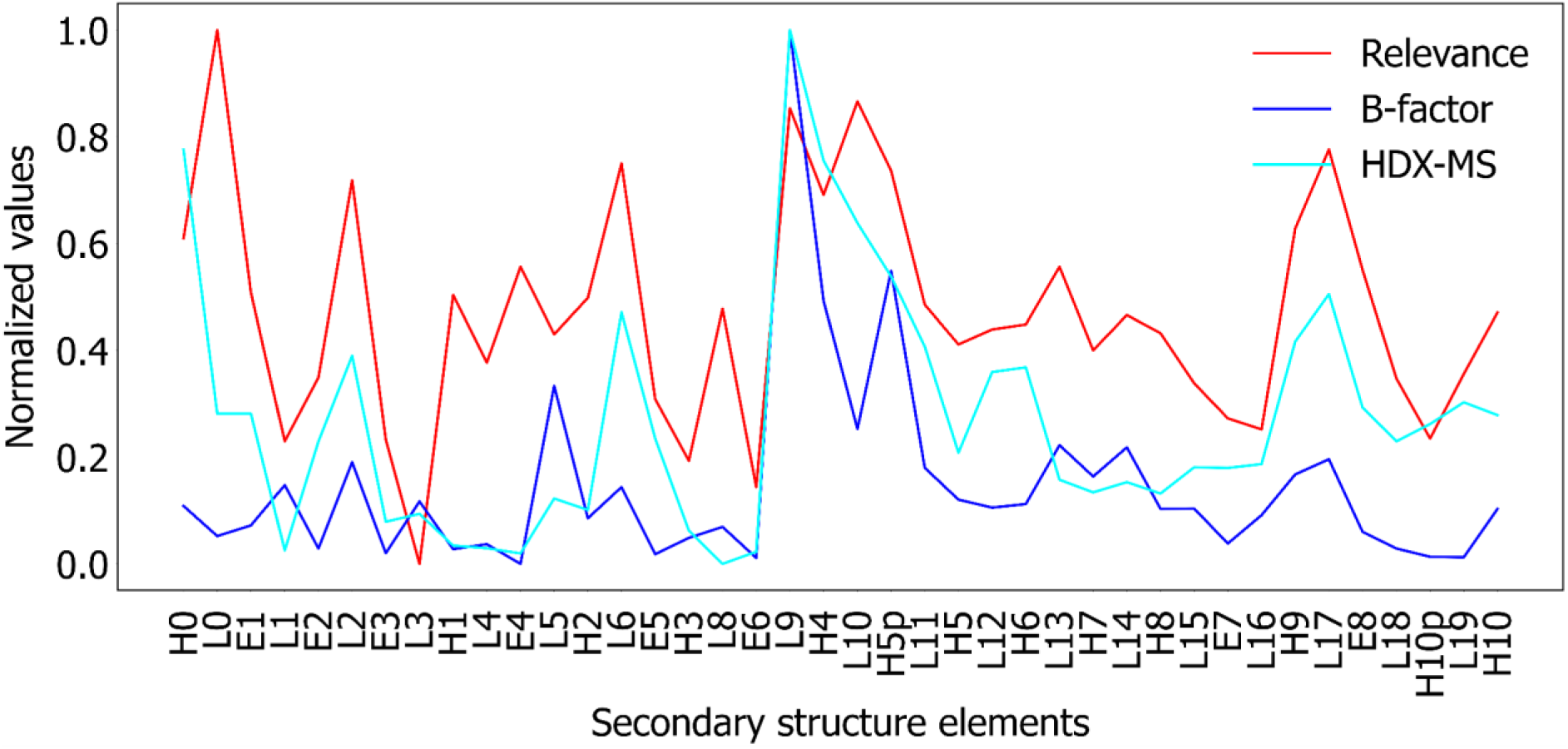
Relevance scores compared with computational B-factors and HDX-MS aggregated per secondary structure element for Anc^HLD-Rluc^. The normalized datasets show moderate-to-strong associations: Pearson correlations of *r* = 0.50 (Relevance vs. B-factor), 0.66 (Relevance vs. HDX-MS), and 0.69 (B-factor vs.HDX-MS); Spearman correlations of ρ = 0.51, 0.62, and 0.49, respectively; and cosine similarities of 0.71, 0.94, and 0.81, respectively.

**Figure S6.**
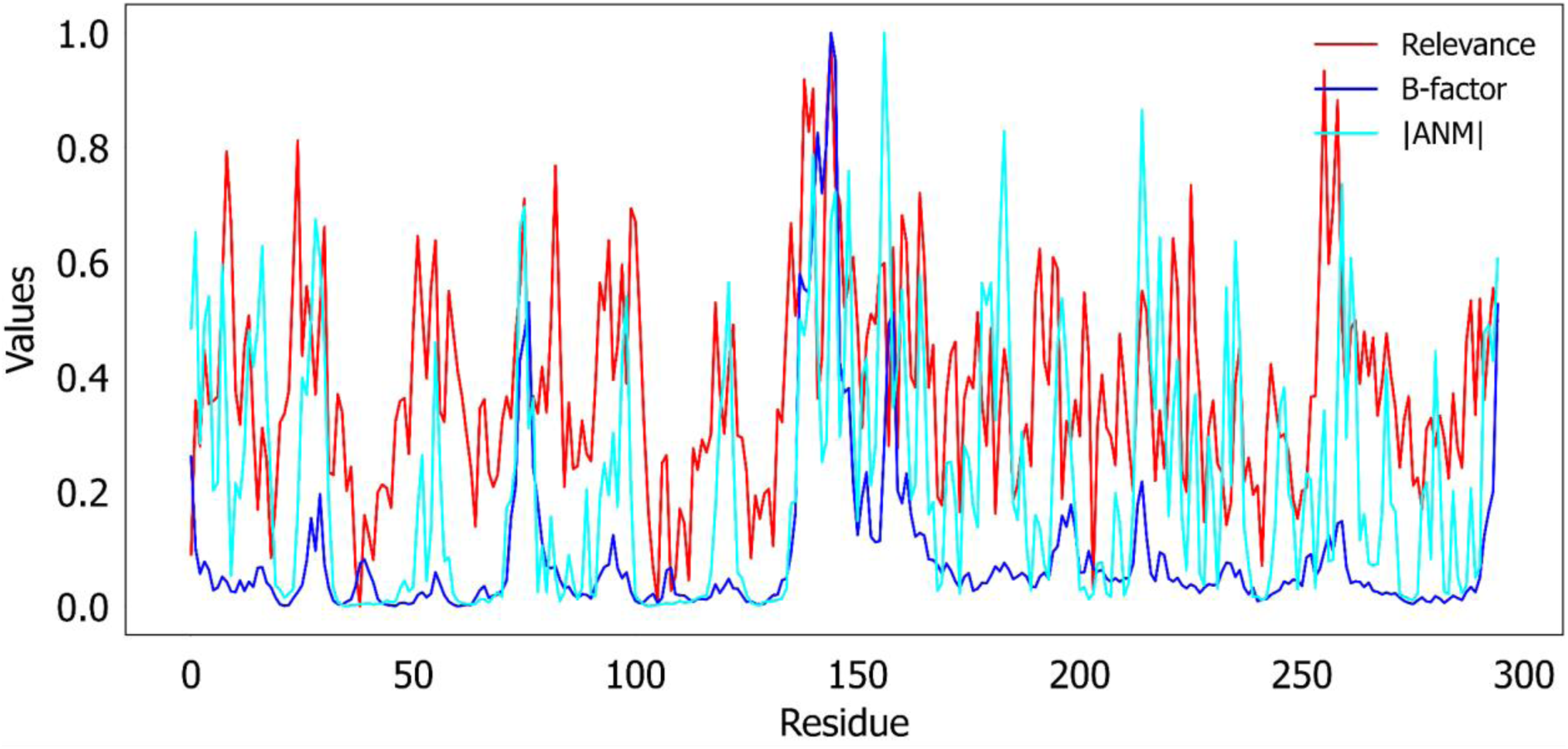
Scaled comparison of Relevance (red), absolute values of ANM (cyan), and B-factors (dark blue) per residue for Anc^HLD-Rluc^. Correlation analysis indicates that Relevance is positively associated with |ANM| (Spearman ρ = 0.52, Pearson r = 0.45, cosine similarity = 0.79) and B-factors (Spearman ρ = 0.42, Pearson r = 0.45, cosine similarity = 0.67). In addition, |ANM| shows a strong positive correlation with B-factors (Spearman ρ = 0.68, Pearson r = 0.52, cosine similarity = 0.71).

**Figure S7.**
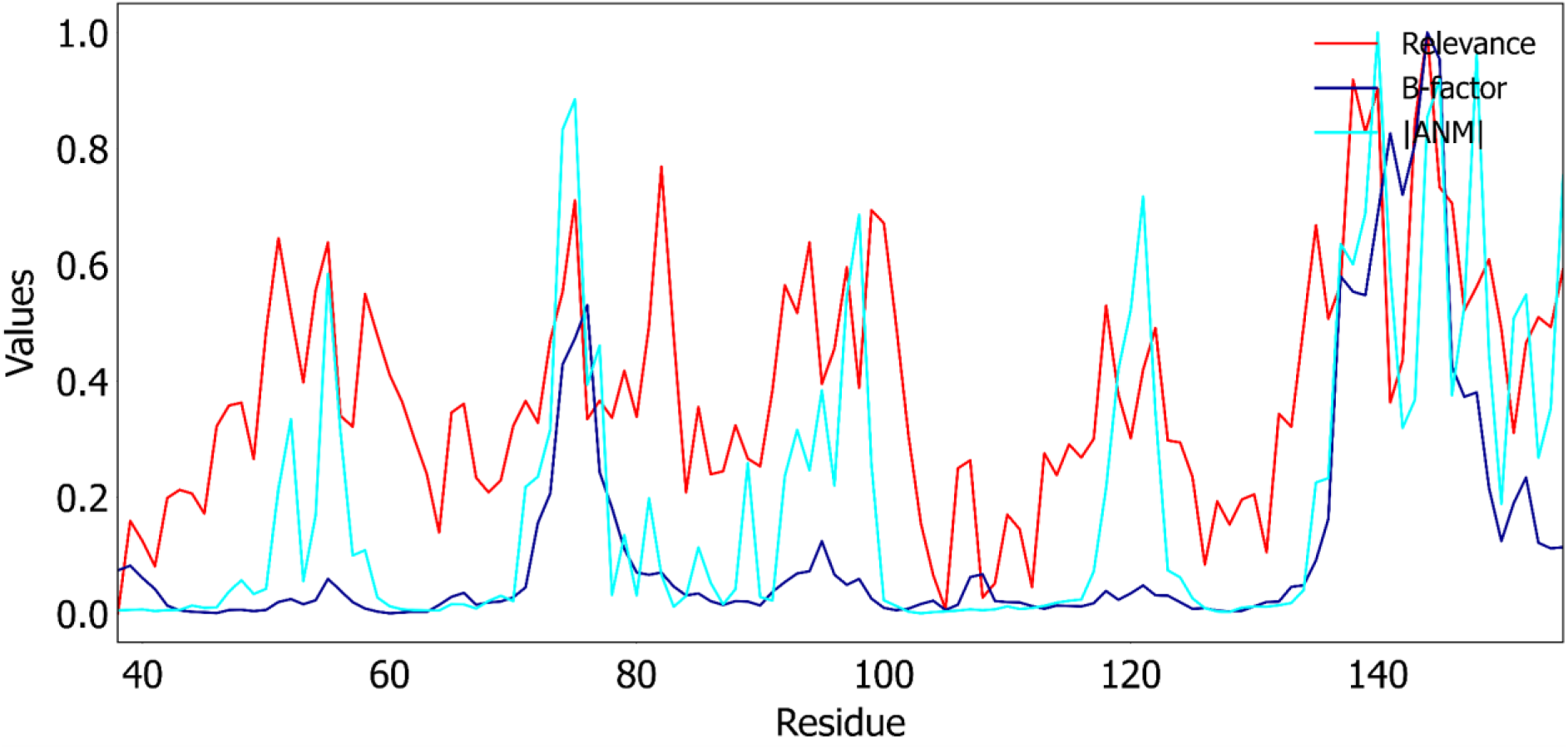
Scaled comparison of Relevance (red), absolute values of ANM (cyan), and B-factors (dark blue) across residues 38–155 for Anc^HLD-Rluc^. Correlation analysis indicates that Relevance is positively associated with |ANM| (Spearman ρ = 0.78, Pearson r = 0.67, cosine similarity = 0.79) and B-factors (Spearman ρ = 0.54, Pearson r = 0.58, cosine similarity = 0.68). In addition, |ANM| shows a strong positive correlation with B-factors (Spearman ρ = 0.70, Pearson r = 0.72, cosine similarity = 0.81).

**Figure S8.**
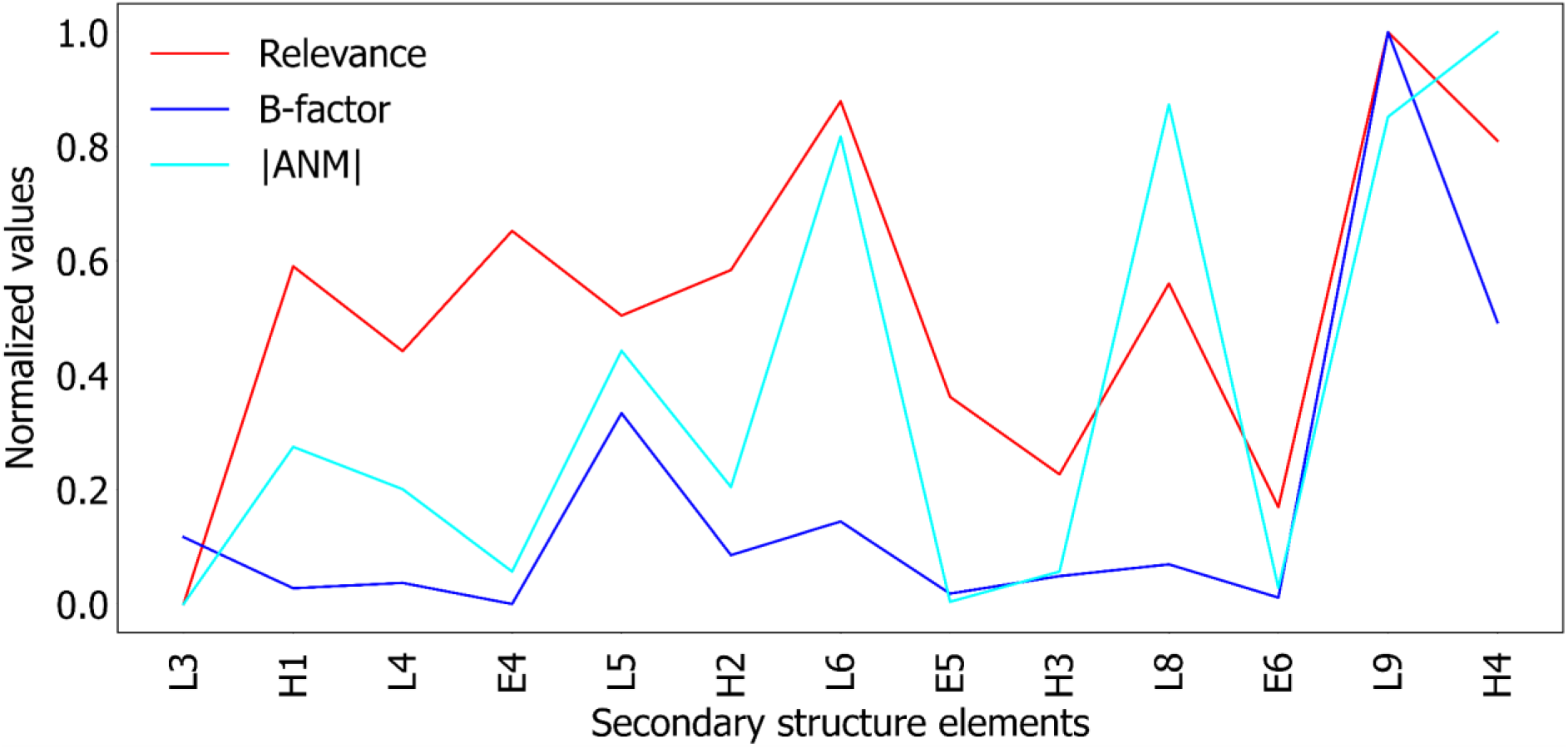
Scaled comparison of Relevance (red), absolute values of ANM (cyan), and B-factors (dark blue) for Anc^HLD-Rluc^, aggregated at the secondary structure level. Correlation analysis indicates that Relevance is positively associated with |ANM| (Spearman ρ = 0.78, Pearson r = 0.77, cosine similarity = 0.86) and B-factors (Spearman ρ = 0.42, Pearson r = 0.61, cosine similarity = 0.74). In addition, |ANM| shows a strong positive correlation with B-factors (Spearman ρ = 0.62, Pearson r = 0.64, cosine similarity = 0.81).

**Figure S9.**
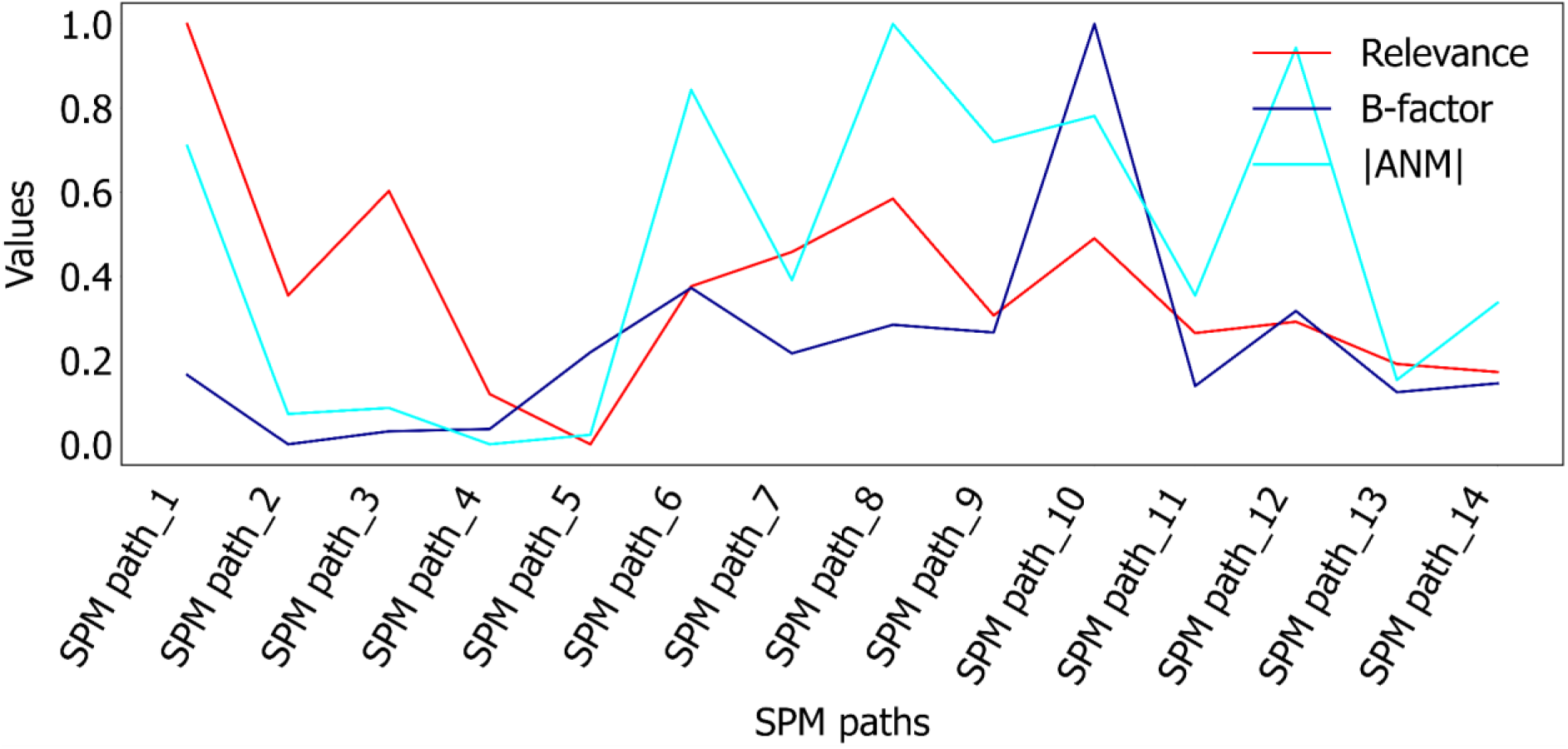
**Scaled comparison of Relevance (red), absolute values of ANM (cyan), and B-factors (dark blue) across shortest-path-map (SPM) pathways for Anc^HLD-RLuc^**. Values are averaged over residues belonging to each SPM path, enabling analysis of dynamical communication routes at the pathway level. Correlation analysis indicates that Relevance is positively associated with |ANM| (Spearman ρ = 0.51, Pearson r = 0.46, cosine similarity = 0.87) and weakly positively associated with B-factors (Spearman ρ = 0.19, Pearson r = 0.14, cosine similarity = 0.81). In addition, |ANM| shows a strong positive correlation with B-factors (Spearman ρ = 0.80, Pearson r = 0.58, cosine similarity = 0.87).

**Figure S10.**
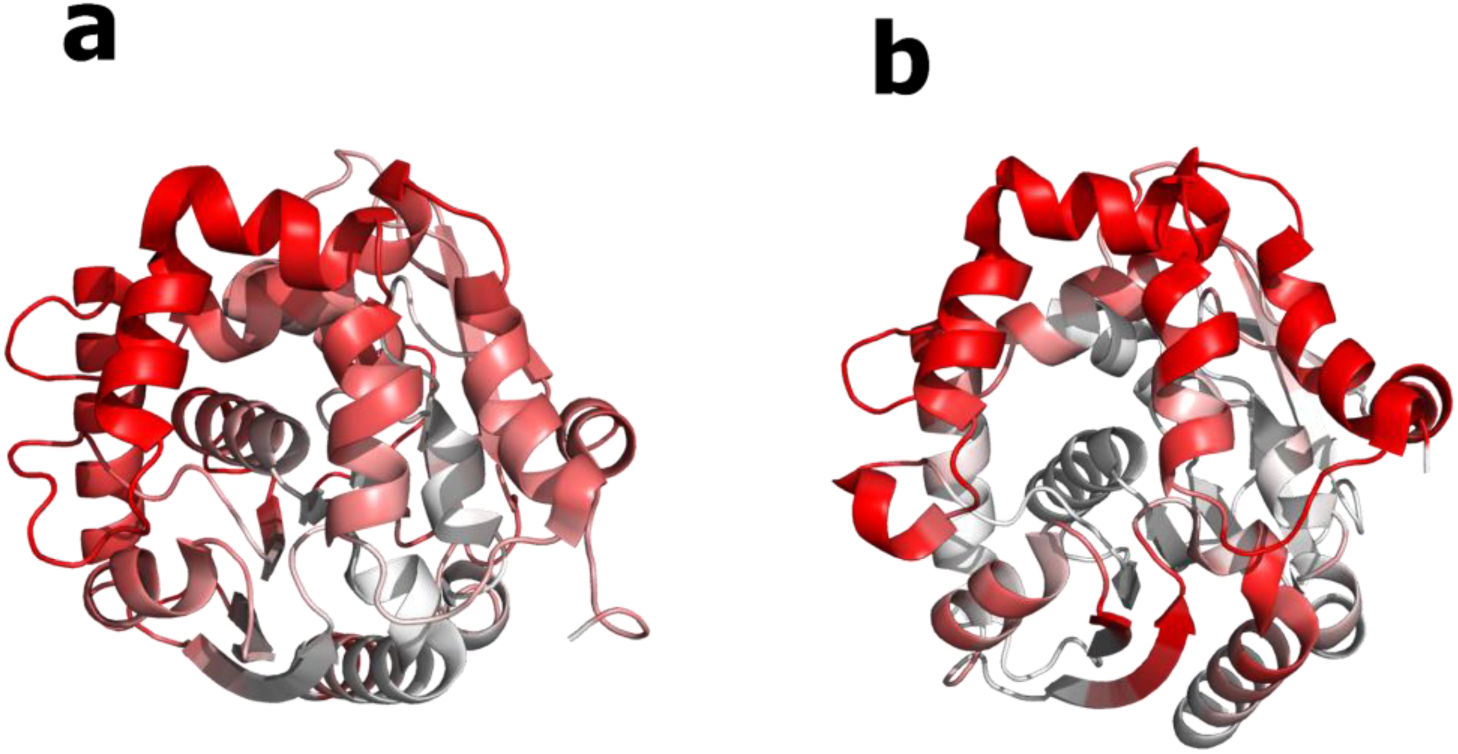
XAI-based dynamic relevance maps for (a) RLuc8 and (b) AncFT. Explainable AI (XAI) analysis applied to MD-derived Cartesian coordinate trajectories to identify residues most influential for predicting conformational evolution in each variant. In panel a (RLuc8), Loop 9 (L9) emerges as the dominant relevance hotspot, consistent with its experimentally established flexibility and central role in cap-domain opening and closing. Additional relevance is observed along helices α4–α5, reflecting the previously reported redistribution of dynamics toward the helix bundle in later-stage luciferase variants. In panel b (AncFT), relevance is similarly concentrated in L9, but with a more pronounced extension into the α4–α5 region and after it compared with Anc^HLD-RLuc^ and RLuc8. This pattern agrees with experimental crystallography, HDX-MS, and MD observations showing that AncFT exhibits intermediate-cap dynamics, with motion partially shifted from L9 toward α5. Together, the XAI results for RLuc8 and AncFT confirm that the model captures the Loop9→α5 redistribution of cap-domain dynamics described in Engineering the protein dynamics of an ancestral luciferase.

**Figure S11.**
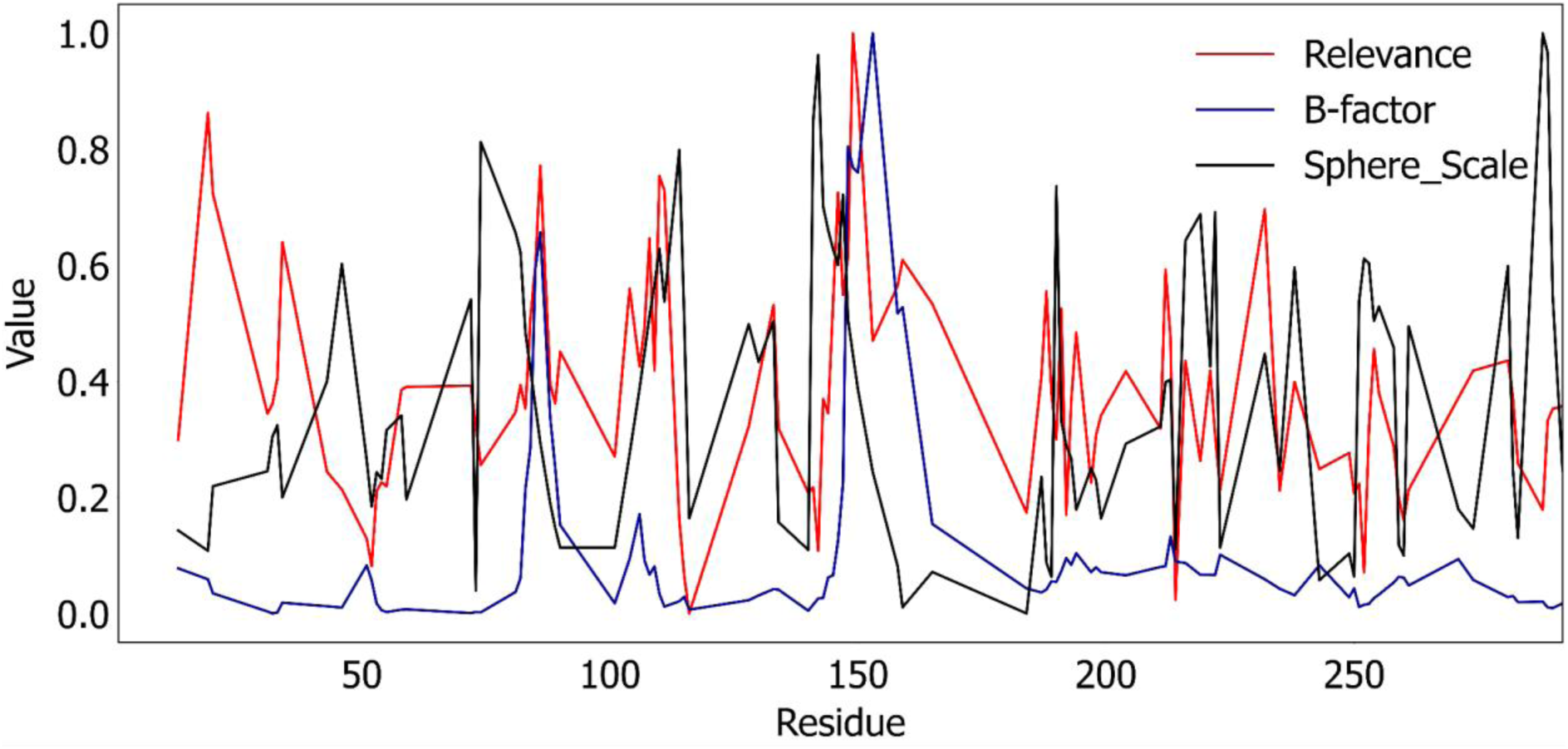
Normalized values of Sphere_Scale (black), relevance values (red), and B-factors (dark blue) per residue for Anc^HLD-RLuc^. Sphere_Scale is a structural descriptor that quantifies the spatial extent of residue motion, reflecting how widely a residue explores its local conformational space. Correlation analysis revealed weak linear and monotonic relationships between Sphere_Scale and relevance values (Spearman: 0.0588, Pearson: 0.0009), but strong similarity in their residue-level spatial patterns as indicated by high cosine similarity (0.9169). Sphere_Scale and B-factors showed weak and slightly negative correlations (Spearman: −0.1763, Pearson: −0.1141) together with moderate cosine similarity (0.5457), suggesting limited correspondence between these descriptors. Relevance values and B-factors exhibited moderate positive correlations (Spearman: 0.4085, Pearson: 0.4933) and relatively high cosine similarity (0.6849), indicating that relevance partially captures experimentally supported flexibility patterns while also reflecting additional structural features associated with collective conformational dynamics.

**Table S1.** Summary of model configurations, data representations, and training strategies.

| System | Data representation | Model architecture | Training setup | Regularization & hyperparameters | Baseline |
| --- | --- | --- | --- | --- | --- |
| **ApoE4** | Ca atoms (residues 24–161); Distance (excluding $i\pm 1$, $i\pm 2$); $\Delta D$ (difference between consecutive frames) | 2D CNN: Conv (4, 8) → Dense (16, 32) → Sigmoid output | Binary classification (apo vs 3-SPA); 80/20 split; 5-fold CV | L2 = 0.1; Dropout = 0.025 (distance), 0.4 ($\Delta D$); LR = $1\times 10^{-4}$ (tested $1\times 10^{-5}$) | Logistic regression (ElasticNet, SAGA) |
| **SAK** | Ca atoms; Distance and $\Delta D$ (excluding $i\pm 1$, $i\pm 2$) | 2D CNN: Conv (2, 4) → Dense (8) → Softmax (4 outputs) | Multiclass classification (4 variants); 80/20 split; 5-fold CV | L2 = 0.1; Dropout = 0.05 | Multinomial logistic regression (ElasticNet, SAGA) |
| **Luciferase** | Ca Cartesian coordinates; next-frame prediction task | 1D CNN autoencoder: 16 → 8 → 3 → 8 → 16 | Supervised reconstruction ($t \rightarrow t+1$); 80/20 split | MSE loss; no scaling; same architecture for all systems | — |

**Table S2.** Summary of model configurations, data representations, and training strategies.

| System | Explainability method | Relevance aggregation | Preprocessing |
| --- | --- | --- | --- |
| **ApoE4** | LRP | Averaged per residue | Subsampling (5 ns), $\Delta D$ computation |
| **SAK** | LRP | Averaged per residue | Subsampling (0.1 ns), $\Delta D$ computation |
| **Luciferase** | LRP | Mean across coordinates and snapshots | Alignment, block-wise subsampling, no scaling |

**Table S3.** Classification performance for ApoE4 using distance-difference (ΔD) representation.

|  | Train |  | 5-fold cross validation |  | Test |  |
| --- | --- | --- | --- | --- | --- | --- |
|  | Accuracy | F1 measure | Accuracy | F1 measure | Accuracy | F1 measure |
| 2D CNN | 0.98 | 0.98 | 0.97 | 0.97 | 0.95 | 0.95 |
| Logistic Regression | 0.54 | 0.68 | 0.49 | 0.64 | 0.45 | 0.60 |

**Table S4.** Residue pairs with the highest LRP relevance scores for each SAK variant.

| Variant | Top five residue pairs with highest relevances |
| --- | --- |
| **42D** | (43, 37), (43, 39), (43, 38), (43, 36), (43, 35), (65, 58) |
| **STAR** | (31, 41), (31, 40), (30, 40), (80, 33), (10, 15) |
| **SY155** | (67, 43), (37, 65), (66, 43), (68, 43), (69, 43) |
| **THR174** | (73, 76), (72, 75), (74, 71), (73, 77), (70, 74) |

**Table S5.** Classification performance for SAK using distance-difference (ΔD) representation.

|  | Train |  | 5-fold cross validation |  | Test |  |
| --- | --- | --- | --- | --- | --- | --- |
|  | Accuracy | F1 measure | Accuracy | F1 measure | Accuracy | F1 measure |
| 2-D CNN | 0.98 | 0.98 | 0.98 | 0.98 | 0.99 | 0.99 |
| Logistic Regression | 0.32 | 0.32 | 0.18 | 0.18 | 0.17 | 0.17 |

**Table S6.** Normalized contributions of Relevance, ANM (absolute values), and B-factor across residue SPM paths for Anc^HLD-Rluc^. Values represent the fraction of each feature’s total (after Min–Max scaling) attributed to individual SPM paths, providing a residue-range–level comparison of structural flexibility (B-factor), dynamics (ANM), and ML relevance.

| Paths | Relevance | ANM | B-factor |
| --- | --- | --- | --- |
| SPM path_1 | 0.127283 | 0.105291 | 0.056281 |
| SPM path_2 | 0.06983 | 0.019615 | 0.02149 |
| SPM path_3 | 0.091961 | 0.021537 | 0.027998 |
| SPM path_4 | 0.048943 | 0.009925 | 0.029144 |
| SPM path_5 | 0.038305 | 0.012946 | 0.067688 |
| SPM path_6 | 0.071765 | 0.123103 | 0.099999 |
| SPM path_7 | 0.079045 | 0.062413 | 0.067119 |
| SPM path_8 | 0.090305 | 0.144161 | 0.081473 |
| SPM path_9 | 0.065585 | 0.106507 | 0.077613 |
| SPM path_10 | 0.081926 | 0.114838 | 0.232339 |
| SPM path_11 | 0.061864 | 0.057471 | 0.050734 |
| SPM path_12 | 0.064262 | 0.136556 | 0.088397 |
| SPM path_13 | 0.055319 | 0.030481 | 0.047647 |
| SPM path_14 | 0.053607 | 0.055156 | 0.052077 |

**Table S7.**
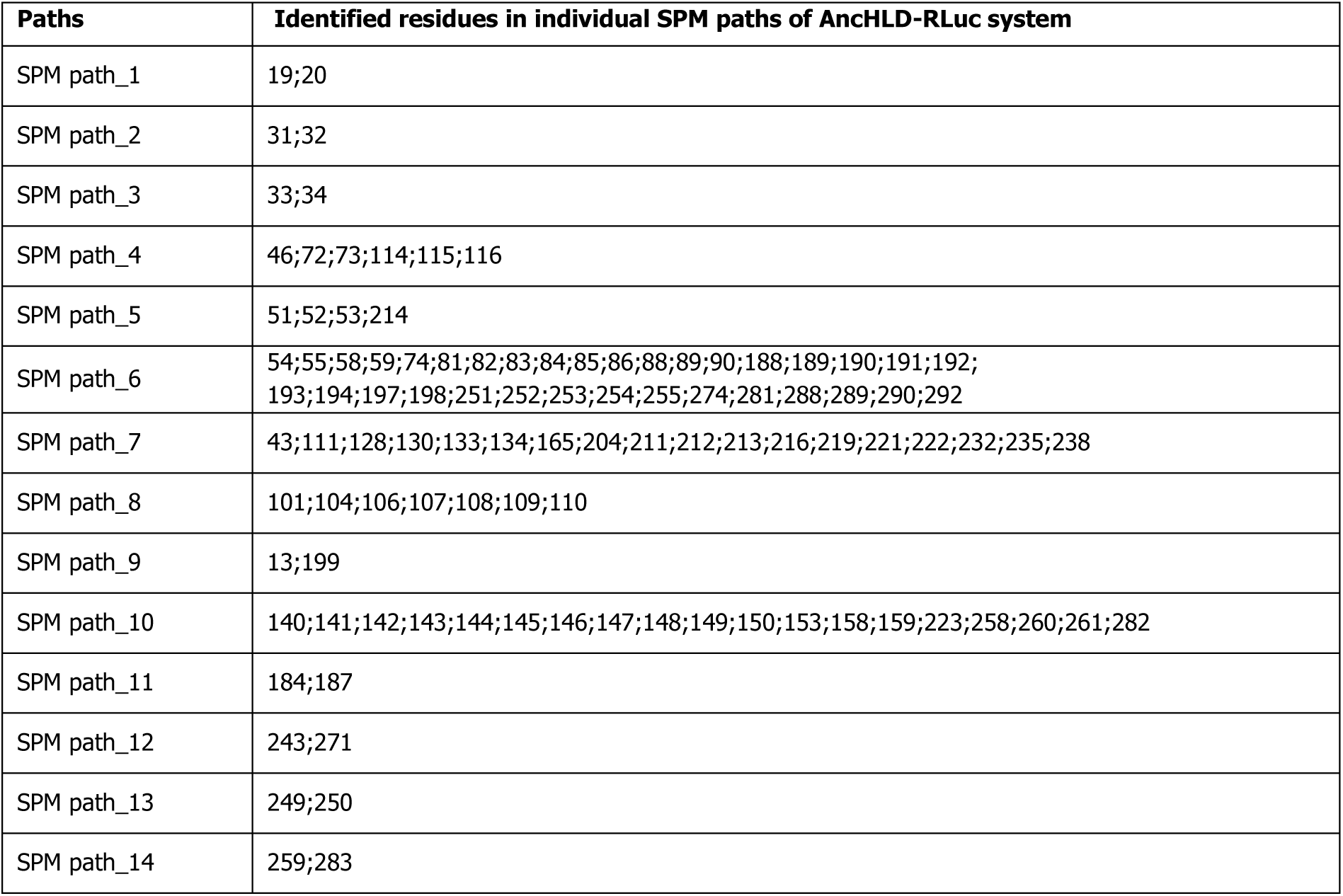
Shortest path map (SPM) method applied for adaptive sampling trajectories of the Anc^HLD-RLuc^ protein. Following the protocol published previously^1^, we identified the SPM paths using a dedicated web server with default hyperparameter settings (Significance - 0.3; Distance - 0.6).

| Paths | Identified residues in individual SPM paths of Anc<sup>HLD-RLuc</sup> system |
| --- | --- |
| SPM path_1 | 19;20 |
| SPM path_2 | 31;32 |
| SPM path_3 | 33;34 |
| SPM path_4 | 46;72;73;114;115;116 |
| SPM path_5 | 51;52;53;214 |
| SPM path_6 | 54;55;58;59;74;81;82;83;84;85;86;88;89;90;188;189;190;191;192;<br>193;194;197;198;251;252;253;254;255;274;281;288;289;290;292 |
| SPM path_7 | 43;111;128;130;133;134;165;204;211;212;213;216;219;221;222;232;235;238 |
| SPM path_8 | 101;104;106;107;108;109;110 |
| SPM path_9 | 13;199 |
| SPM path_10 | 140;141;142;143;144;145;146;147;148;149;150;153;158;159;223;258;260;261;282 |
| SPM path_11 | 184;187 |
| SPM path_12 | 243;271 |
| SPM path_13 | 249;250 |
| SPM path_14 | 259;283 |

<sup>1</sup> Casadevall, G.; Casadevall, J.; Duran, C.; Osuna, S. The Shortest Path Method (SPM) Webserver for Computational Enzyme Design. Protein Engineering, Design and Selection 2024, 37, gzae005. https://doi.org/10.1093/protein/gzae005.

